# A deep learning framework for inference of single-trial neural population dynamics from calcium imaging with sub-frame temporal resolution

**DOI:** 10.1101/2021.11.21.469441

**Authors:** Feng Zhu, Harrison A. Grier, Raghav Tandon, Changjia Cai, Anjali Agarwal, Andrea Giovannucci, Matthew T. Kaufman, Chethan Pandarinath

## Abstract

In many brain areas, neural populations act as a coordinated network whose state is tied to behavior on a moment-by-moment basis and millisecond timescale. Two-photon (2p) calcium imaging is a powerful tool to probe network-scale computation, as it can measure the activity of many individual neurons, monitor multiple cortical layers simultaneously, and sample from identified cell types. However, estimating network state and dynamics from 2p measurements has proven challenging because of noise, inherent nonlinearities, and limitations on temporal resolution. Here we describe RADICaL, a deep learning method to overcome these limitations at the population level. RADICaL extends methods that exploit dynamics in spiking activity for application to deconvolved calcium signals, whose statistics and temporal dynamics are quite distinct from electrophysiologically-recorded spikes. It incorporates a novel network training strategy that capitalizes on the timing of 2p sampling to recover network dynamics with high temporal precision. In synthetic tests, RADICaL infers network state more accurately than previous methods, particularly for high-frequency components. In real 2p recordings from sensorimotor areas in mice performing a “water grab” task, RADICaL infers network state with close correspondence to single-trial variations in behavior, and maintains high-quality inference even when neuronal populations are substantially reduced.

## Introduction

In recent years, advances in neural recording technologies have enabled simultaneous monitoring of the activity of large neural populations^1–3^. These technologies are enabling new insights into how neural populations implement the computations necessary for motor, sensory, and cognitive processes^4^. However, different recording technologies impose distinct tradeoffs in the types of questions that may be asked^5–7^. Modern electrophysiology enables access to hundreds to thousands of neurons within and across brain areas with high temporal fidelity^2^. Yet in any given area, electrophysiology is limited to a sparse sampling of relatively active, unidentified neurons^6^ (**Fig. 1a**). In contrast, two photon (2p) calcium imaging offers the ability to monitor the activity of vast populations of neurons - rapidly increasing from tens of thousands to millions^3,8,9^ - in 3-D, often with identified layers and cell types of interest^10,11^. Thus 2p imaging is a powerful tool for understanding how neural circuitry gives rise to function.

**Figure 1.**
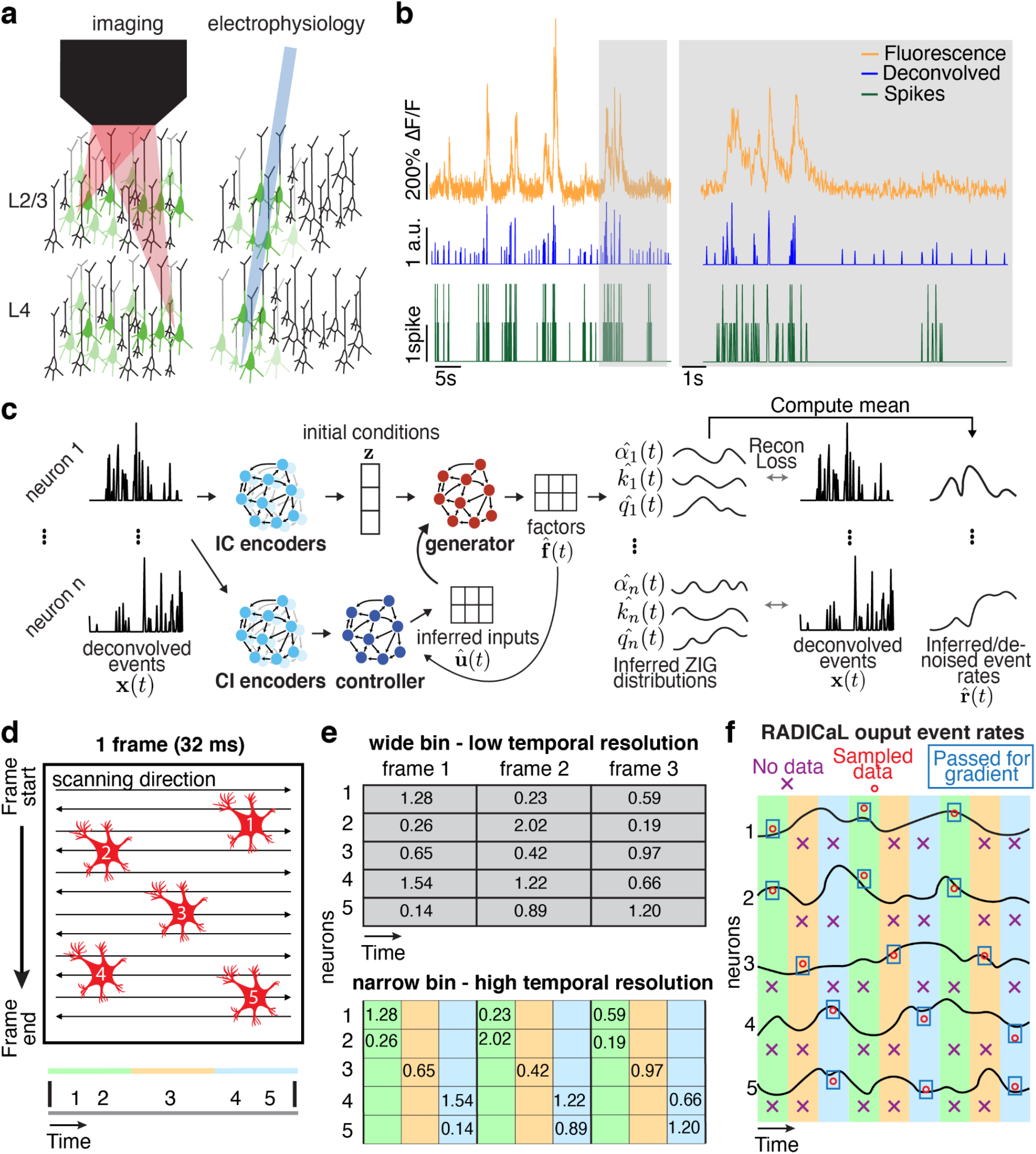
Improving inference of network state from 2p imaging. (a) Calcium imaging offers the ability to monitor the activity of many neurons simultaneously, in 3-D, often with cell types of interest and layers identified. In contrast, electrophysiology sparsely samples the neurons in the vicinity of a recording electrode, and may be biased toward neurons with high firing rates. (b) Calcium fluorescence transients are a low-passed and lossy transformation of the underlying spiking activity. Spike inference methods may provide a reasonable estimate of neurons’ activity on coarse timescales (*left*), but yield poor estimates on fine timescales (*right*; data from ref. ^7^). (c) RADICaL uses a recurrent neural network-based generative model to infer network state - *i.e*., de-noised event rates for the population of neurons - and assumes a time-varying ZIG observation model. (d) *Top*: in 2p imaging, the laser’s serial scanning results in different neurons being sampled at different times within the frame. *Bottom*: individual neurons’ sampling times are known with sub-frame precision (colors) but are typically analyzed with whole-frame precision (gray). (e) Sub-frame binning precisely captures individual neurons’ sampling times but results in neuron-time points without data. The numbers in the table indicate the deconvolved event in each frame. (f) SBTT is a novel network training method for sparsely sampled data that prevents unsampled time-neuron data points from affecting the gradient computation.

A key tradeoff, however, is that the fluorescence transients measured via calcium imaging are a low-passed and nonlinearly-distorted transformation of the underlying spiking activity (**Fig. 1b**). Further, because neurons are serially scanned by a laser that traverses the field of view (FOV), a trade-off exists between the size of the FOV (and hence the number of neurons monitored), the sampling frequency, and the pixel size (and therefore the signal-to-noise with which each neuron is sampled). These factors together limit the fidelity with which the activity of large neuronal populations can be monitored and extracted via 2p, and thus limit our ability to link activity measured with 2p imaging to neural computation and behavior on fine timescales.

In recent years, a large amount of effort has been dedicated to improving the inference of spike trains from 2p calcium data by detecting calcium influx events, *i.e*., time points where single spikes, or multiple spikes in close succession, produce detectable fluorescence transients^12^. Ideally the spikes-to-fluorescence transformation would be invertible, such that analyzing calcium events would be equivalent to analyzing spiking activity^5^. However, recent benchmarks illustrate that a variety of algorithms to infer calcium events reach a similar ceiling of performance and make consistent predictions, and all achieve limited correspondence to ground truth spiking activity obtained with electrophysiology, particularly on fine timescales^13,14^.

Rather than focusing on the responses of individual neurons, an alternative approach is to characterize patterns of covariation across a neuronal population to reveal the multi-dimensional internal state of the network as a whole. These “latent variable models”, or simply “latent models”, describe each neuron’s activity as a reflection of the whole network’s state over time. For example, when applied to electrophysiological data, latent models assume that an individual neuron’s spiking is a noisy observation of a latent “firing rate”, which fluctuates in a coordinated way with the firing rates of other neurons in the population. Despite their abstract nature, the trajectory of network state inferred by latent models can reveal key insights into the computations being performed by the brain areas of interest^4^. Inferred network state can also enhance our ability to relate neural activity to behavior. For example, one state-of-the-art deep learning method to estimate network state from electrophysiological spiking data is Latent Factor Analysis via Dynamical Systems (LFADS)^15,16^. In applications to data from motor, sensory, and cognitive regions, LFADS reveals rules that govern how network state evolves over time consistently across behavioral conditions, while also revealing tight correspondences with single-trial behavior on a 5-10 millisecond timescale^16,17^.

Given the success of latent models in uncovering network state from electrophysiological data, here we develop a new approach to achieve accurate inference of network state from activity monitored through 2p calcium imaging. We first begin with LFADS, and evaluate network state inference using simulated 2p data in which activity reflects known, nonlinear dynamical systems, and with real 2p data from mice performing a water reaching task. LFADS uncovers network state with substantially higher accuracy then standard approaches (e.g., deconvolution plus Gaussian smoothing). We then develop the Recurrent Autoencoder for Discovering Imaged Calcium Latents (RADICaL) to improve inference over LFADS through innovations tailored specifically for 2p data. In particular, we modify the network architecture to better account for the statistics of deconvolved calcium signals, and develop a novel network training strategy that exploits the staggered timing of 2p sampling of neuronal populations to achieve precise, sub-frame temporal resolution. Our new approach substantially improves inference of network state from 2p data, shown in synthetic data through accurate recovery of high-frequency features (up to 20 Hz), and in real data through improved prediction of neuronal activity, as well as prediction of single-trial variability in hand kinematics during rapid reaches (lasting 200-300 ms). Ultimately, RADICaL provides an avenue to tie precise, population-level descriptions of neural computation with the anatomical and circuit details revealed via calcium imaging.

## Results

### Leveraging population dynamics to infer network state from 2p imaging data

Dynamical systems models such as LFADS rely on two key principles to infer network state from neural population activity. First, simultaneously recorded neurons exhibit coordinated spatial patterns of activation that reflect the state of the network^18,19^. Due to this coordination, network state might be reliably estimated even if the measurement of individual neurons’ activity is unreliable. Second, these coordinated spatial patterns evolve over time based on consistent rules (dynamics)^4,20^. Thus, while it may be challenging to accurately estimate the network’s state based on activity at a single time point, knowledge of the network’s dynamics provides further information to help constrain network state estimates using data from multiple time points.

To apply these principles to improve inference from 2p data, we extended LFADS to produce RADICaL (**Fig. 1c**). Both LFADS and RADICaL model neural population dynamics using recurrent neural networks (RNNs) in a sequential autoencoder configuration (details in *Methods*, and in previous work^15,16^). This configuration is built on the assumption that the network state underlying neural population activity can be approximated by an input-driven dynamical system, and that observed activity is a noisy observation of the state of the dynamical system. The dynamical system itself is modeled by an RNN (the ‘generator’). For any given trial, the time-varying network state can be captured by three pieces of information: the initial state of the dynamical system (trial-specific), the dynamical rules that govern state evolution (shared across trials), and any time-varying external inputs that may affect the dynamics (trial-specific). The states of the generator are linearly mapped onto a latent space to produce a ‘factors’ representation, which is then transformed to produce the time-varying output for each neuron (detailed below). The model has a variety of hyperparameters that control training and prevent overfitting, whose optimal settings are not known *a priori*. To ensure that these hyperparameters were optimized properly for each dataset, we built RADICaL on top of a powerful, large-scale hyperparameter optimization framework we recently developed known as AutoLFADS^17,21^.

### Novel features of RADICaL

RADICaL incorporates two major innovations over LFADS and AutoLFADS. First, we modified RADICaL’s observation model to better account for the statistics of deconvolved events. In LFADS, discrete spike count data are modeled as samples from an underlying time-varying Poisson process for each neuron. However, deconvolving 2p calcium signals results in a time series of continuous-valued events, with imperfect correspondence to the actual spike times and counts^13^. These deconvolved events can be better approximated at each timepoint by a zero-inflated gamma (ZIG) distribution, which combines a gamma distribution to model the calcium event magnitudes and a point mass that represents the elevated probability of zero values^22^. In RADICaL, deconvolved events are therefore modeled as samples from a time-varying ZIG distribution whose parameters are taken from the output of the generator RNN (**Fig. 1c**; details in *Methods*). We define the network state at any given time point as a vector containing the inferred (i.e., de-noised) event rates of all neurons, where the de-noised event rate is taken as the mean of each neuron’s inferred ZIG distribution at each time point (equation (3) in *Methods*). The de-noised event rates are latent variables that are tied to the underlying network state at each time point. In LFADS, the instantaneous intensity parameter of the Poisson process completely specifies the spike count distribution for a neuron, while in RADICaL, the ZIG distribution requires three parameters (detailed in *Methods*). The RADICaL generator RNN must therefore produce multiple parameters that do not directly correspond to the biological network’s activity. To avoid this complication, rather than using the factors themselves as an estimate of the biological network’s state, we used the de-noised event rates. Doing so for both RADICaL and AutoLFADS allowed us to compare methods as directly as possible.

Second, we developed a novel neural network training strategy, selective backpropagation through time^23^ (SBTT), that leverages the precise sampling times of individual neurons to enable recovery of high-frequency network dynamics. Since standard 2p microscopes rely on point-by-point raster scanning of a laser beam to acquire frames, it is possible to determine the sample times for each neuron with high precision within the frame (**Fig. 1d**). To leverage this information to improve inference of high-frequency network dynamics on single trials, we recast the underlying interpolation problem as a missing data problem: we treat imaging a whole frame as sequentially imaging multiple, smaller bands containing different neurons. In this framing, each neuron is effectively sampled sparsely in time, *i.e*., the majority of time points for each neuron do not contain valid data (**Fig. 1e**). Such sparsely sampled data creates a challenge when training the underlying neural network: briefly, neural networks are trained by adjusting their parameters (weights), and performing this adjustment requires evaluating the gradient of a cost function with respect to weights. SBTT allows us to compute this gradient using only the valid data, and ignore the missing samples (**Fig. 1f**; see *Methods*). Because SBTT only affects how we compute the gradient and update the weights, the network still infers event rates for every neuron at every time point, regardless of whether samples exist at that time point or not. This allows the trained network to accept sparsely-sampled observations as input, and produce high-temporal resolution event rate estimates as its output.

### RADICaL uncovers high-frequency features from simulated data

We first tested RADICaL using simulated 2p data, which provides a valuable tool for quantifying performance because the underlying network state is known and parameterizable. We hypothesized that the new features of RADICaL would allow it to infer higher-frequency features with greater accuracy than standard approaches, such as Gaussian-smoothing the deconvolved events (“s-deconv”), smoothing the simulated fluorescence traces themselves (“s-sim-fluor”), or state-of-the-art tools for electrophysiology analysis, such as AutoLFADS. We generated synthetic spike trains by simulating a population of neurons whose firing rates were linked to the state of a Lorenz system^15,24^ (detailed in *Methods* and **Supp. Fig. 1a**). We ran the Lorenz system at various speeds, allowing us to investigate the effects of temporal frequency on the quality of network state recovery achieved by different methods. In the 3-dimensional Lorenz system, the *Z* dimension contains the highest-frequency content (**Supp. Fig. 1b**). Here we denote the frequency of each Lorenz simulation by the peak frequency of the power spectrum of its *Z* dimension (**Supp. Fig. 1c**).

We used the synthetic spike trains to generate realistic noisy fluorescence signals consistent with GCAMP6f (detailed in *Methods* and **Supp. Fig. 2**). To recreate the variability in sampling times due to 2p laser scanning, fluorescence traces were simulated at 100 Hz and then sub-sampled at 33.3 Hz, with offsets in each neuron’s sampling times consistent with spatial distributions across a simulated FOV. We then deconvolved the generated fluorescence signals to extract events ^25,26^. Because RADICaL uses SBTT, it could be applied directly to the deconvolved events with offset sampling times. In contrast, for both AutoLFADS and s-deconv, deconvolved events for all neurons were treated as all having the same sampling times (i.e., consistent with the frame times), as is standard in 2p imaging (detailed in *Methods*).

Despite the distortions introduced by the fluorescence simulation and deconvolution process, RADICaL was able to infer event rates that closely resembled the true underlying rates (**Fig. 2a**). To assess whether each method accurately inferred the time-varying state of the Lorenz system, we mapped the representations from the different approaches - i.e., the event rates inferred by RADICaL or AutoLFADS, the smoothed deconvolved events, and the smoothed simulated fluorescence traces - onto the true underlying Lorenz states using cross-validated ridge regression. We then quantified performance using the coefficient of determination (*R^2^*), which quantifies the fraction of the variance of the true latent variables captured by the estimates. **Figure 2b** shows the Lorenz *Z* dimension for example trials from three Lorenz speeds, as well as the recovered values for three of the methods. RADICaL inferred latent states with high fidelity (*R^2^*>0.8) up to 15 Hz, and significantly outperformed other methods across a range of frequencies (**Fig. 2c**; performance for the *X* and *Y* dimensions is shown in **Supp. Fig. 3**; p<0.05 for all frequencies and dimensions, paired, one-sided t-Test, detailed in *Methods*). Notably, performance in estimating latent states was improved due to both of the innovations in RADICaL (i.e., modeling events with a ZIG distribution, and SBTT), with SBTT contributing more (**Supp. Fig. 4**). To test RADICaL’s ability in estimating single-trial dynamics for a task that lacks a repetitive trial-structure, we varied the simulation so that each trial has a unique initial condition for the Lorenz system. RADICaL accurately inferred the latent states on single trials (**Supp. Fig. 5a**) and outperforms AutoLFADS and s-deconv at high Lorenz oscillation frequencies (**Supp. Fig. 5b**).

**Figure 2.**
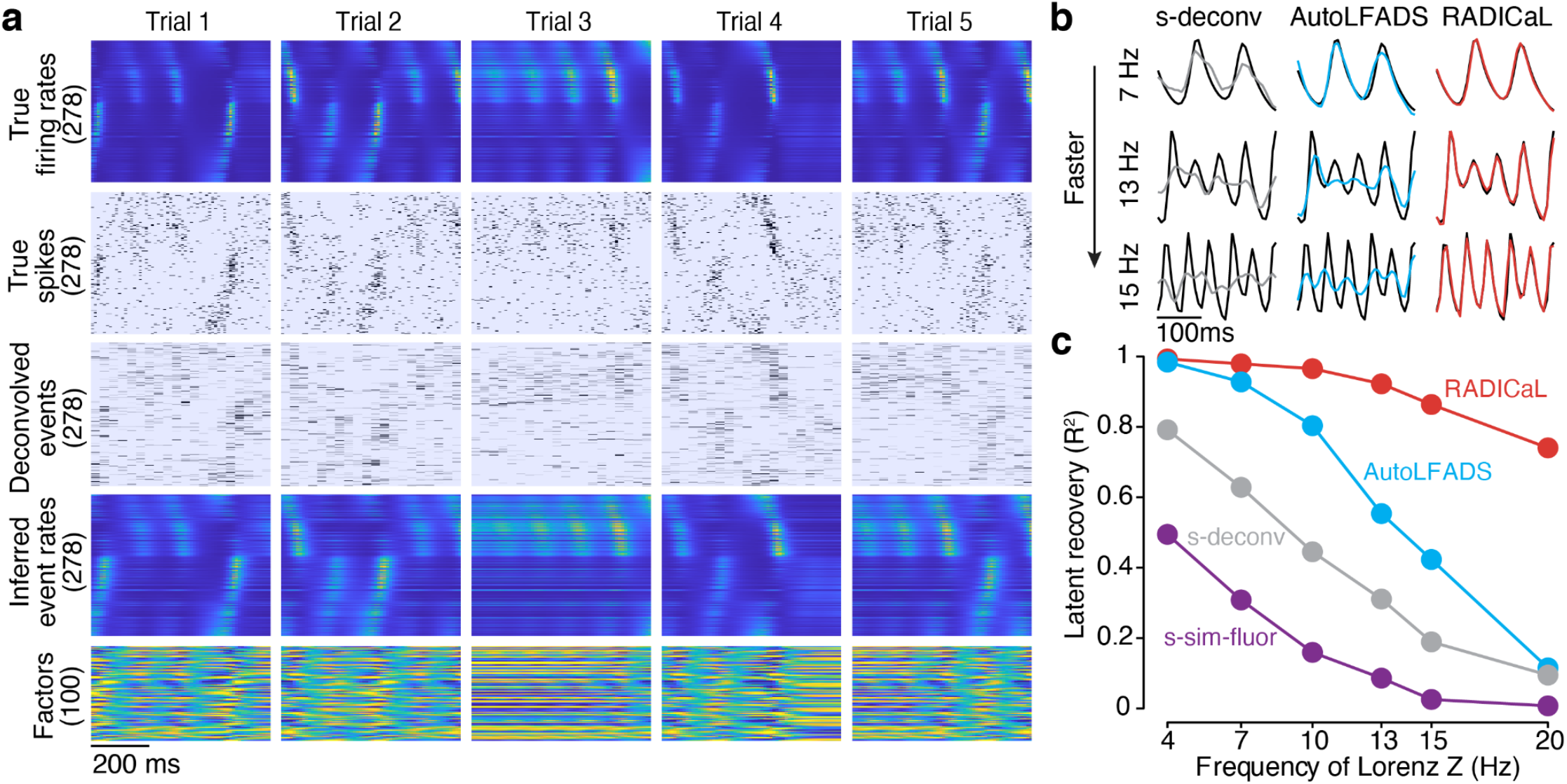
Application of RADICaL to synthetic data. (a) Example firing rates and spiking activity from a Lorenz system simulated at 7 Hz, deconvolved calcium events (inputs to RADICaL), and the corresponding rates and factors inferred by RADICaL. Simulation parameters were tuned so that the performance in inferring spikes using OASIS matched previous benchmarks^13^ (see *Methods*). (b) True and inferred Lorenz latent states (*Z* dimension) for a single example trial from Lorenz systems simulated at three different frequencies. Black: true. Colored: inferred. (c) Performance in estimating the Lorenz *Z* dimension as a function of simulation frequency was quantified by variance explained (*R^2^*) for all 4 methods.

These synthetic results provide an important proof-of-principle that RADICaL can infer high-frequency features of the network activity underlying 2p signals, which is readily validated when ground truth is known. To better understand the regimes in which RADICaL will recover the underlying latent variables well or poorly, we performed variants of the simulation experiments along 4 additional axes: imaging speed (**Supp. Fig. 6**), high frequency structure in the latent variables (**Supp. Fig. 7**), noise levels (**Supp. Fig. 8**), and whether RADICaL could be effective when used with algorithms that infer spike times instead of event rates, such as MLspike^27^ (**Supp. Fig. 9**). In all cases we found that RADICaL substantially outperformed alternate approaches. However, as expected, our analysis showed that deconvolution itself performs poorly at very slow sampling rates (e.g., 2Hz and below), and for very high frequency content (e.g., >20 Hz), and thus RADICaL’s performance in those regimes is limited by the use of deconvolution as a preprocessing step.

These additional tests demonstrate RADICaL’s ability to extend into different use cases. However, it is important to acknowledge limitations of the simulation process that might constrain the generality of these results when applied to real data. In particular, the parameter space of possible experiments is very large, especially considering the variety of calcium indicators, protein expression patterns, imaging settings, cell types and firing rate patterns. An exhaustive search of this parameter space is infeasible and thus it is difficult to know whether the results from any particular choice of simulation parameters (or a variety of choices) can be extrapolated to real experimental conditions. Thus, we next benchmarked performance on real data to demonstrate RADICaL’s utility in the real world.

### RADICaL improves network state inference in data from a mouse water grab task

We next tested RADICaL on 2p recordings from mice performing a forelimb water grab task (**Fig. 3a**, *top*). We analyzed data from four experiments: two mice with two sessions from each mouse, in which different brain areas were imaged (M1, S1). Our task was a variant of the water-reaching task of Galiñanes & Huber^28^. In each trial, the mouse was cued by the pitch of an auditory tone to reach to a left or right spout and retrieve a droplet of water with its right forepaw (**Fig. 3a**, *bottom*; see *Methods*). The forepaw position was tracked at 150 frames per second with DeepLabCut^29^ for 420-560 trials per experiment. To test whether each method could reveal structure in the neural activity at finer resolution than left vs. right reaches, we divided trials from each condition into subgroups based on forepaw height during the reach (**Fig. 3a**, *top right*; see *Methods*). Two-photon calcium imaging from GCaMP6f transgenic mice was performed at 31 Hz, with 430-543 neurons within the FOV in each experiment (**Fig. 3b**).

**Figure 3.**
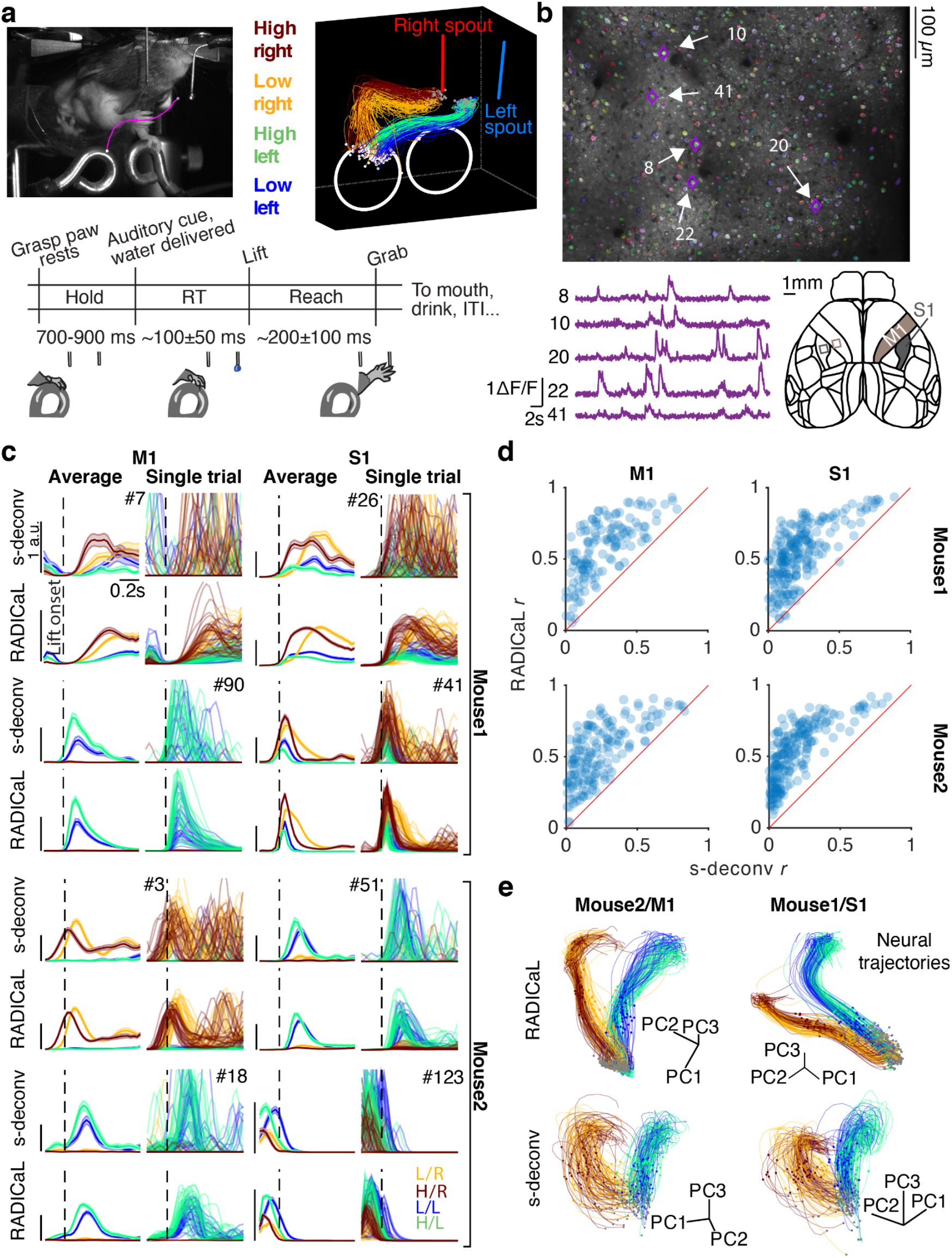
Application of RADICaL to real two-photon calcium imaging of a water grab task. (a) Task. *Top left*: Mouse performing the water grab task. Pink trace shows paw centroid trajectory. Bottom: Event sequence/task timing. RT: reaction time. ITI: inter-trial interval. *Top right*: Individual reaches colored by subgroup identity. (b) *Top*: an example field of view (FOV), identified neurons colored randomly. *Bottom left*: dF/F from a single trial for 5 example neurons. *Bottom right*: Allen Atlas M1/S1 brain regions imaged. (c) Comparison of trial-averaged (left) and single-trial (right) rates for 8 individual neurons for two different brain areas (left vs. right) and two different mice (top half vs. bottom half) for s-deconv and RADICaL (alternating rows). *Left*: each trace represents a different reach subgroup (4 in total) with error bars indicating s.e.m. *Right*: each trace represents an individual trial (same color scheme as trial-averaged panels). *Odd rows*: s-deconv event rates (Gaussian kernel: 40 ms s.d.). *Even rows*: RADICaL-inferred event rates. Horizontal scale bar represents 200 ms. Vertical scale bar denotes event rate (a.u.). Vertical dashed line denotes lift onset time. (d) Performance of RADICaL and s-deconv in capturing the empirical PSTHs on single trials. Correlation coefficient *r* was computed between the inferred single-trial event rates and empirical PSTHs. Each point represents an individual neuron. (e) Single-trial neural trajectories derived from RADICaL rates (top row) and s-deconv rates (bottom row) for two experiments (*left*: Mouse2/M1; *right*: Mouse1/S1), colored by subgroups. Each trajectory is an individual trial, plotting from 200 ms before to 400 ms after lift onset. Lift onset times are indicated by the dots in the same colors with the trajectories. Grey dots indicate 200 ms prior to lift onset time. Neural trajectories from additional experiments are shown in **Supp. Fig. 11**.

With real datasets, a key challenge when benchmarking latent variable inference is the lack of ground truth data for comparison. A useful first-order assessment is whether the event rates inferred for individual trials match the empirical peri-stimulus time histograms (PSTHs), *i.e*., the rates computed by averaging noisy single-trial data across trials with similar behavioral characteristics^16,17^. While this approach obscures meaningful across-trial variability, it provides a ‘denoised’ estimate that is useful for coarse performance quantification and comparisons. To compute empirical PSTHs, we averaged the smoothed deconvolved events (s-deconv rates) across trials within each subgroup.

We found that RADICaL-inferred event rates recapitulated features of individual neurons’ activity that were apparent in the empirical PSTHs, both when averaging across trials, but also on individual trials (**Fig. 3c**). Importantly, RADICaL is an unsupervised method, meaning that it was not provided any behavioral information, such as whether the mouse reached to the left or right on a given trial, or which subgroup a trial fell into. Yet the single-trial event rates inferred by RADICaL showed clear separation not only between left and right reach conditions, but also between subgroups of trials within each condition. This separation was not clear with the single-trial s-deconv rates. We quantified the correspondence between the single-trial inferred event rates and the empirical PSTHs via Pearson’s correlation coefficient (*r*; see *Methods*). RADICaL single-trial event rates showed substantially higher correlation with the empirical PSTHs than s-deconv rates (**Fig. 3d**) or those inferred by AutoLFADS (**Supp. Fig. 10**). Importantly, these improvements were not limited to a handful of neurons, but instead were broadly distributed across the population.

We next tested whether the population activity inferred by RADICaL also showed meaningful structure on individual trials. We produced low-dimensional visualizations of the population’s activity by applying principal component analysis (PCA) to the RADICaL-inferred or s-deconv event rates after log-transforming and trial-averaging, and then projected the single-trial event rates (also log-transformed) into the subspace formed by the top three PCs. The low-D trajectories computed from the RADICaL-inferred rates showed consistent, clear single-trial structure that corresponded to behavioral conditions and subgroups for all four experiments (**Fig. 3e**, *top row*; **Supp. Fig. 11**, *top row*), despite RADICaL receiving no direct information about which trials belonged to which condition. In comparison, low-D trajectories computed from the s-deconv rates showed noisy single-trial structure with little correspondence to behavioral subgroups (**Fig. 3e**, *bottom row*; **Supp. Fig. 11**, *bottom row*). To provide a quantitative summary, we measured the distance of the low-D trajectories between each trial and other trials across subgroups (d_across_) vs. within the same subgroup (d_within_) for any given time and computed the distance ratio (detailed in *Methods*). The distance ratio (i.e., d_across_ / d_within_) of RADICaL-derived trajectories was higher than s-deconv-derived trajectories across time points, which was also consistent across four experiments (**Supp. Fig. 12**). To further test whether RADICaL could pick up on idiosyncrasies of single-trial activity, we evaluated a subset of trials that were “loopy” and highly non-stereotyped, showing multiple large peaks in hand velocity (**Supp. Fig. 13a**). RADICaL captured the distinct patterns of neural responses for these loopy trials, without receiving any behavioral information (**Supp. Fig. 13b**).

### RADICaL captures dynamics that improve behavioral prediction

We next tested whether the RADICaL-inferred event rates were closely linked to behavior by decoding forepaw positions and velocities from the inferred event rates using cross-validated ridge regression (**Fig. 4a**). Decoding using RADICaL-inferred rates significantly outperformed results from s-deconv rates, or from the AutoLFADS-inferred rates (**Fig. 4b**; position: average *R*^2^ of 0.91 across all experiments, versus 0.75 and 0.85 for s-deconv and AutoLFADS, respectively; velocity: average *R*^2^ of 0.62 across the mice/areas, versus 0.37 and 0.51 for s-deconv and AutoLFADS, respectively; p<0.05 for position and velocity for all individual experiments, paired, one-sided t-test, detailed in *Methods*). Improvements achieved by RADICaL were shown on most trials (**Supp. Fig. 15**). Importantly, the performance advantage was not achieved by simply predicting the mean event rates for all trials of a given condition: RADICaL also outperformed AutoLFADS and s-deconv in decoding the kinematic residuals (i.e., the single-trial deviations from the mean; **Supp. Fig. 16**). To assess how decoding improvements were distributed as a function of frequency, we computed the coherence between the true and decoded positions and velocities for each method (**Fig. 4c**). RADICaL predictions showed higher coherence with behavior than predictions from s-deconv or AutoLFADS across a wide range of frequencies, and the difference in coherence between RADICaL and AutoLFADS widened (especially for position) at higher frequencies (5-15 Hz). This argues that RADICaL improved decoding particularly because it improved recovery of higher-frequency features of the neural activity. Notably, decoding was improved due to both innovations in RADICaL (i.e., modeling events with a ZIG distribution, and SBTT), and the combination of the two innovations significantly improved performance over each innovation alone (**Supp. Fig. 17**).

**Figure 4.**
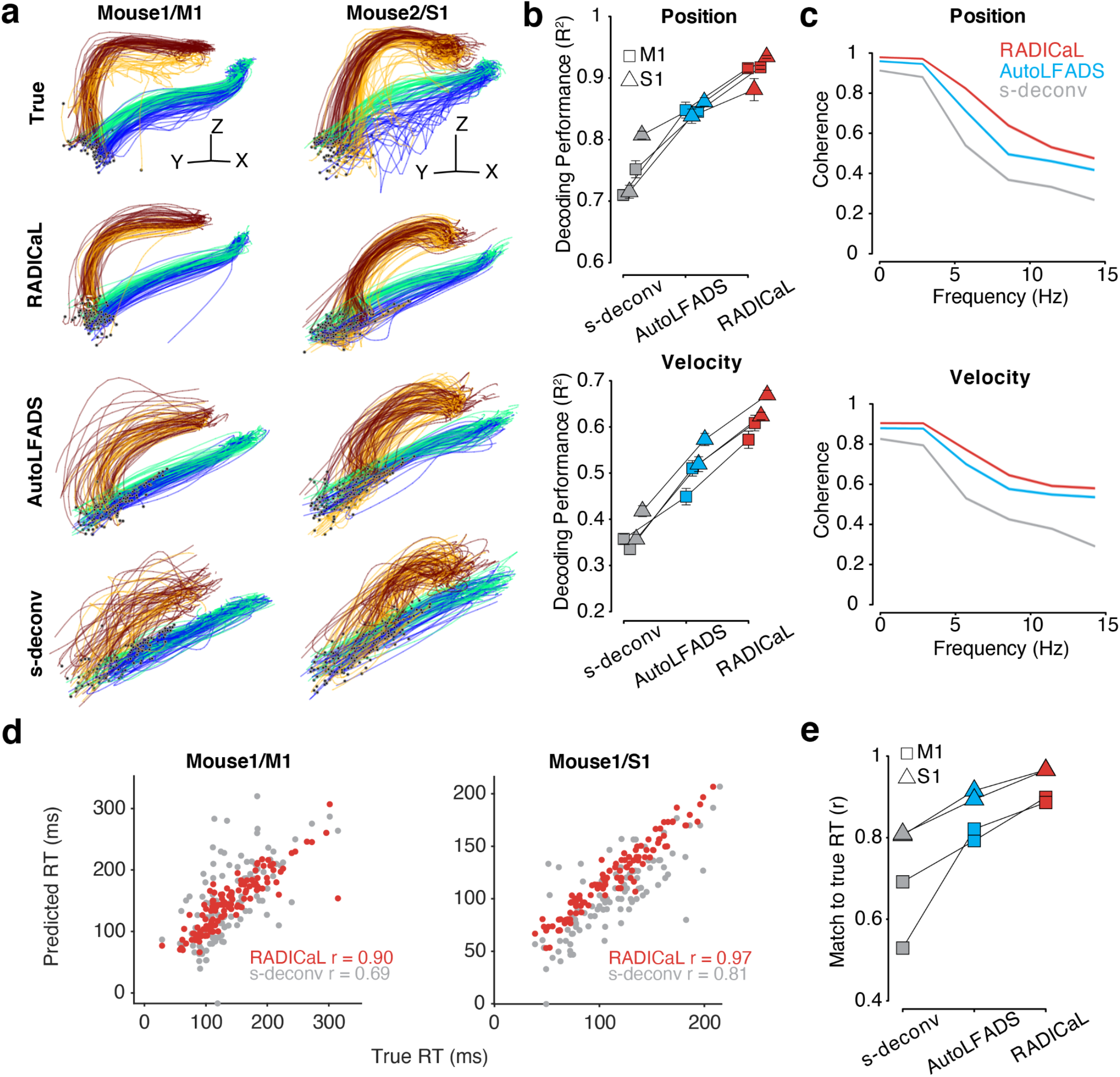
RADICaL improves prediction of behavior. (a) Decoding hand kinematics using ridge regression. Each column shows an example mouse/area. Row 1: true hand positions trajectories, colored by subgroups. Rows 2–4: predicted hand positions using ridge regression applied to the event rates inferred by RADICaL or AutoLFADS, or s-deconv rates (Gaussian kernel: 40 ms s.d.). Hand positions from additional experiments are shown in **Supp. Fig. 14**. (b) Decoding accuracy was quantified by measuring variance explained (*R*^2^) between the true and decoded position (top) and velocity (bottom) across all trials across each of the 4 datasets (2 mice for M1, denoted by squares, and 2 mice for S1, denoted by triangles), for all 3 techniques. (c) Quality of reconstructing the kinematics across frequencies was quantified by measuring coherence between the true and decoded position (top) and velocity (bottom) for individual trials across all 4 datasets, for all 3 techniques. (d) Predicting single-trial reaction times using RADICaL or s-deconv rates. Each dot represents an individual trial, color-coded by event rate inference method. Correlation coefficient *r* was computed between the true and predicted reaction times. Prediction of single-trial reaction times from additional experiments are shown in **Supp. Fig. 18**. (e) Performance of predicting single-trial reaction times across each of the 4 datasets (2 mice for M1, denoted by squares, and 2 mice for S1, denoted by triangles), for all 3 techniques.

We next tested whether RADICaL could capture meaningful trial-to-trial variability by predicting reaction time (RT) from the inferred event rates using cross-validated logistic regression^30^ (detailed in *Methods*). The RT in a trial is defined as the time between water presentation and movement onset. RTs predicted from RADICaL-inferred rates showed high correlation with the true RTs (**Fig. 4d**), and outperformed results from s-deconv rates, or from the AutoLFADS-inferred rates (**Fig. 4e**; average *r* of 0.93 across all experiments, versus 0.71 and 0.86 for s-deconv and AutoLFADS, respectively).

### RADICaL retains high decoding performance when reducing the number of neurons used in the model

In previous demonstrations on electrophysiological spiking data, LFADS maintained accurate performance in reconstructing single-trial neural activity and decoding even when reducing the number of sampled neurons^16^. To evaluate whether this holds for RADICaL, we performed a neuron-downsampling experiment where we gradually reduced the number of neurons used in training RADICaL or AutoLFADS, either in a random fashion (**Fig. 5**), or in a FOV-shrinking fashion (**Supp. Fig. 19**). In both cases, RADICaL retained relatively high decoding performance as the population size was reduced. Decoding performance declined gradually, with a steeper slope for velocity. Notably, however, performance when only 25% of the neurons were used for training RADICaL was similar to that of AutoLFADS - and higher than for s-deconv - when those methods were applied to the full population of neurons. Analyses were robust to the seed used for selecting different random subsets of neurons (**Supp. Fig. 20**). These results provide an avenue to retain information when scanning sparser populations (such as when a cell type of interest is in the minority), smaller areas when imaging deep structures with a limited FOV due to a GRIN lens, or using smaller FOVs to capture multiple layers or regions while retaining overall frame rate (see *Discussion*).

**Figure 5.**
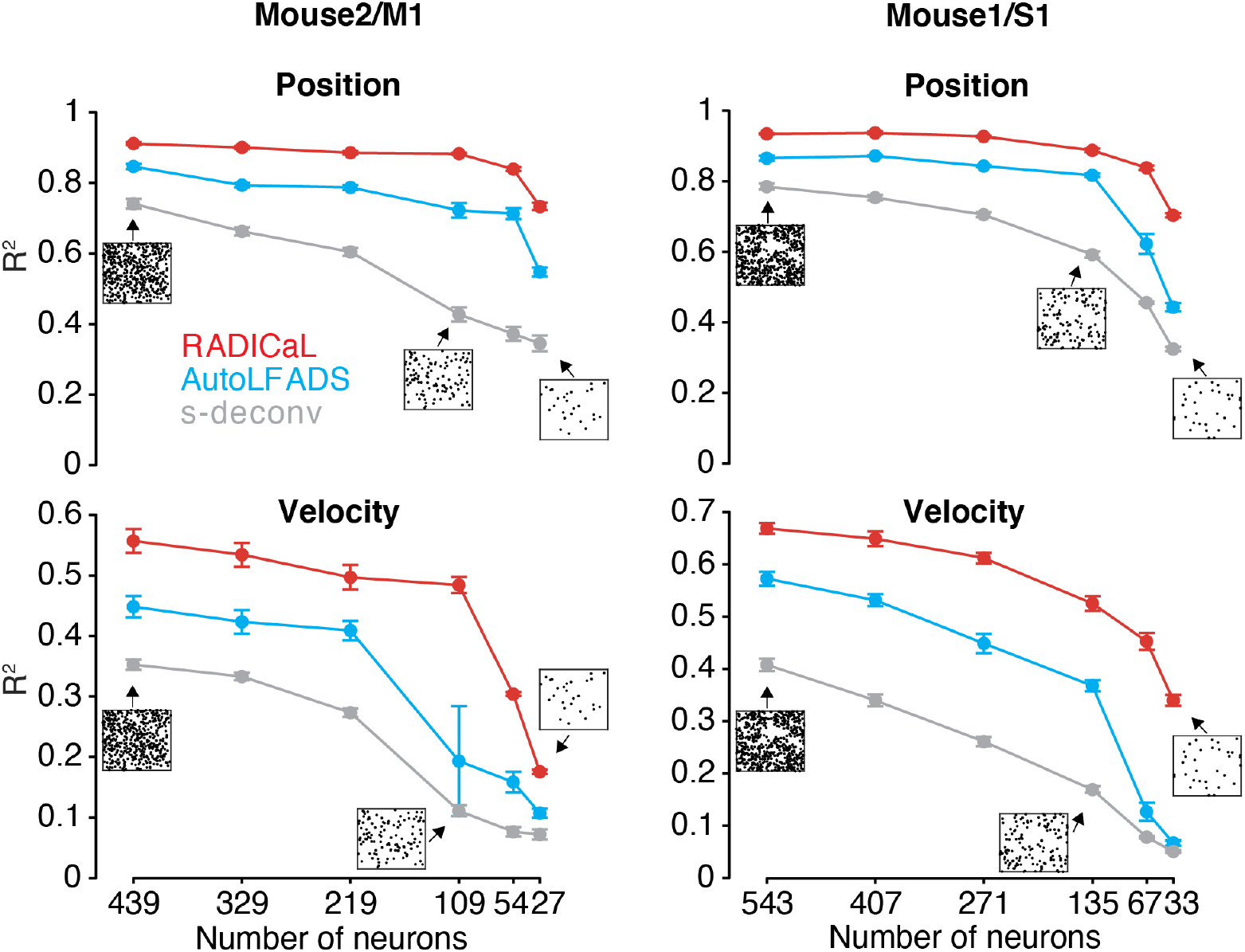
RADICaL retains high decoding performance in a neuron downsampling experiment. Decoding performance was measured as a function of the number of neurons used in each technique (top: Position; bottom: Velocity). Data are from Mouse2/M1 (left) and Mouse1/S1 (right). Performance was quantified using variance explained (*R^2^*). Figure insets indicate the selected neurons in the FOV for the full population of neurons and examples for different subsets. Each black dot in the insets represents a neuron.

## Discussion

2p imaging is a widely-used method for interrogating neural circuits, with the potential to monitor vast volumes of neurons and provide new circuit insights that elude electrophysiology. To date, however, it has proven challenging to precisely infer network state from imaging data, due in large part to the inherent noise, indicator dynamics, and low temporal resolution associated with 2p imaging. RADICaL bridges this gap. RADICaL is tailored specifically for 2p imaging, with a noise emissions model that is appropriate for deconvolved calcium events, and a novel network training strategy (SBTT) that takes advantage of the specifics of 2p laser scanning to achieve substantially higher temporal resolution. Through synthetic tests, we demonstrated that RADICaL accurately infers network state and substantially outperforms alternate approaches in uncovering high-frequency fluctuations. Then, through careful validation on real 2p data, we demonstrated that RADICaL infers network state trajectories that are closely linked to single-trial behavioral variability, even on fast timescales. Finally, we demonstrated that RADICaL maintains high-quality inference of network state even as the neural population size is reduced substantially.

The ability to de-noise neural activity on single trials is highly valuable. First, de-noising improves the ability to decode behavioral information from neural activity, allowing subtle relationships between neural activity and behavior to be revealed (**Fig. 4**). Second, de-noising may enable the field to move away from experimental paradigms that evoke stereotyped behaviors needed to facilitate trial-averaging of neural data. This support for reduced stereotypy could allow greater insight in experiments with animals such as mouse and marmoset, where powerful experimental tools are available but highly repeatable behaviors are challenging to achieve. A move away from trial-averaging could also enable better interpretability of more complex or naturalistic behaviors^17,31–34^. Third, this de-noising capability will enable greater insight into processes that fundamentally differ from trial to trial, such as learning from errors^35,36^, variation in internal states such as arousal^37,38^, or paradigms in which tuning to uninstructed movements contaminates measurement of the task-related behavioral variables of interest^39^. Finally, this de-noising greatly improves inference of network state (**Fig. 2**), mitigating some of the known distortions of neural activity introduced by calcium imaging^5^. Importantly, both of the widely used modalities for recording populations of neurons with single-cell resolution (electrophysiology and calcium imaging) have distinct advantages and disadvantages, and both provide biased information about the underlying neural population^6^. Whereas LFADS has served as a powerful tool for denoising electrophysiology data and accurately inferring network state, no similar method existed for the complementary technique of calcium imaging; RADICaL fills this gap.

In recent years, a variety of computational methods have been developed to analyze 2p imaging data^12^. 2p preprocessing pipelines^8,26^ normally include methods that correct for brain motion, localize and demix neurons’ fluorescence signals, and infer event rates from fluorescence traces. Several studies have applied deep learning in attempts to improve spike inference^40–42^, while a few others have focused on uncovering population-level structure^43–48^ or locally linear dynamics underlying population activity, in particular via switching linear dynamical systems-based methods^49,50^. Here we built RADICaL on the AutoLFADS architecture, which leverages deep learning and large-scale distributed training. This enables the integration of more accurate observation models (ZIG) and powerful optimization strategies (SBTT), while potentially inheriting the high performance and generalized applicability previously demonstrated for AutoLFADS^17^.

Many behaviors are performed on fast timescales (e.g., saccades, reaches, movement correction, etc.), and thus previous work has made steps in overcoming the limits of modest 2p frame rates in attempts to infer the fast changes in neural firing rates that relate to these fast behaviors. Efforts to chip away at this barrier have relied on regularities imposed by repeated stimuli or highly stereotyped behavior^51,52^, or jittered inferred events on sub-frame timescales to minimize the reconstruction error of the associated fluorescence^40^. RADICaL takes a different approach. In particular, it links sub-frame timing to neural population dynamics, representing a more powerful and generalizable approach that does not require stereotypy in the behavior or neural response and which could therefore be applied to datasets with more naturalistic or flexible behaviors.

Though we made an effort to test with realistic simulations, investigating various settings along multiple axes, and on real 2p data from both M1 and S1, it remains untested how RADICaL would generalize to other experimental settings. Noise levels can span a wide range in real experiments, depending on the optics, calcium indicators, expression levels, and other factors. Behaviors can vary in complexity and population dynamics can be high-dimensional. Though it is not guaranteed that RADICaL would work in all possible settings, it provides a solution to the spatiotemporal tradeoff that is inherent to any scanning technique, which enables retaining temporal resolution while increasing the spatial area of sampling.

As shown in our simulated experiments, deconvolution places an upper bound on RADICaL’s performance, limiting its potential in slow sampling regimes (i.e., 2 Hz) with fast indicators or in more challenging inference cases (e.g., higher-frequency latent content, higher noise levels, etc). Because deconvolution outputs an event rate for each frame - which is a summary of the cumulative effect of the spikes within the frame - it necessarily discards some high-frequency features in the data. To mitigate these limitations, future work could build an end-to-end model that integrates the generative rates- to-fluorescence process and operates on the fluorescence traces directly. Complementary work has begun exploring in this direction^53^, but our unique innovation of selective backprop through time presents an opportunity to greatly improve the quality of recovering high-frequency features when the sampling rate is limited. More broadly, as benchmarking efforts are an invaluable resource for systematically comparing methods and building on advances from various different developers^54^, carefully-designed benchmarking efforts for network state inference from 2p data could accelerate progress in this field.

The ability to achieve high-quality network state inference despite limited neuronal population size opens the door to testing new choices about how to perform the experiments themselves. For example, it could enable understanding the role of an uncommon neuronal subtype, or the single-trial outputs of an area by imaging projection neurons that are sparsely distributed throughout that area. With subcortical structures that require relay lenses, it could extract more information from a smaller FOV, permitting the use of a smaller relay lens that causes less damage to overlying brain structures. Or, when hopping between different layers^10,11^ or brain areas^55,56^, fewer lines could be imaged per FOV to retain a higher overall frame rate while achieving good inference from each FOV. When the number of neurons within each FOV is limited, one further advantage that RADICaL inherits from LFADS is that it allows for multi-session stitching^16^, which could provide an avenue to combine data from different sessions to improve inference of the underlying dynamics for each FOV.

In sum, RADICaL provides a framework to push back the limits of the space-time tradeoff in 2p calcium imaging, enabling accurate inference of population dynamics in vast populations and with identified neurons. Future work will explore how best to exploit these capabilities for different experimental paradigms, and to link the power of dynamics with the anatomical detail revealed with calcium imaging.

## Acknowledgements

We thank M. Rivers and R. Vescovi for help with the high-speed camera setup, D. Sabatini for contributions to the behavioral control software, and T. Abe and A. Mosberger for help adapting RADICaL for NeuroCAAS. This work was supported by the Emory Neuromodulation and Technology Innovation Center (ENTICe), NSF NCS 1835364, NIH Eunice Kennedy Shriver NICHD K12HD073945, the Simons Foundation as part of the Simons-Emory International Consortium on Motor Control, NIH NINDS/OD DP2 NS127291, NIH BRAIN/NIDA RF1 DA055667 (CP), the Alfred P. Sloan Foundation (CP, MTK), NSF NCS 1835390, The University of Chicago, the Neuroscience Institute at The University of Chicago (MTK), NIH NINDS R01 NS121535 (MK), and a Beckman Young Investigators Award (AG). The work was also supported by the following collaborative awards (PI: Prof. Ellen Hess, Emory): NIH NINDS R21 NS116311, Imagine, Innovate and Impact (I3) Funds from the Emory School of Medicine and through the Georgia CTSA NIH UL1-TR002378, and a pilot grant from the Emory Udall Center of Excellence for Parkinson’s Research.

## Code availability

RADICaL for Google Cloud Platform can be downloaded from GitHub at github.com/snel-repo/autolfads and the tutorial is available at snel-repo.github.io/autolfads. RADICaL for NeuroCAAS^57^ is available at http://www.neurocaas.org/analysis/17. Source code of RADICaL is available at https://github.com/snel-repo/lfads-cd/tree/radical.

## Data availability

Data will be made available upon reasonable request at the time of publication.

## Author Contributions

F.Z. and C.P. designed the study, with input from A.G. and M.K.. C.P. and M.K. conceptualized the SBTT approach. F.Z. and C.P. performed analyses and wrote the manuscript with input from all other authors. F.Z. and C.P. developed the algorithmic approach. F.Z., C.C., and A.G. developed the simulation pipeline. H.G. and M.K. designed and performed experiments with mice, and developed the real data preprocessing pipeline with input from F.Z. and C.P.. R.T. contributed to initial simulations and data analysis. F.Z., A.G., M.K., and C.P. edited and revised the manuscript with input from all other authors. F.Z. and A.A. adapted RADICaL for Google Cloud Platform and NeuroCAAS.

## Methods

### AutoLFADS and RADICaL architecture and training

The core model that AutoLFADS and RADICaL build on is LFADS. A detailed overview of the LFADS model is given in refs. ^15,16^. Briefly, LFADS is a sequential application of a variational auto-encoder (VAE). A pair of bidirectional RNNs (the initial condition and controller input encoders) operate on the spike sequence and produce initial conditions for the generator RNN and time-varying inputs for the controller RNN. All RNNs were implemented using gated recurrent unit (GRU) cells. At each time step, the generator state evolves with input from the controller and the controller receives delayed feedback from the generator. The generator states are linearly mapped to factors, which are mapped to the firing rate of the neurons using a linear mapping followed by an exponential nonlinearity. The optimization objective is to maximize a lower bound on the likelihood of the observed spiking activity given the rates produced by the generator network, and includes KL and L2 regularization penalties. During training, network weights are optimized using stochastic gradient descent and backpropagation through time.

Identical network sizes were used for both AutoLFADS and RADICaL runs and for both simulation and real 2P data. The dimension of initial condition encoder, controller input encoder, and controller RNNs was 64. The dimension of the generator RNN was 100. The generator was provided with 64-dimensional initial conditions and 2-dimensional controller outputs (i.e., inferred inputs u(t)) and linearly mapped to 100-dimensional factors. The initial condition prior distribution was Gaussian with a trainable mean that was initialized to 0 and a variance that was fixed to 0.1. The minimum allowable variance of the initial condition posterior distribution was set to 1e-4. The controller output prior was autoregressive with a trainable autocorrelation tau and noise variance, initialized to 10 and 0.1, respectively. The Adam optimizer (epsilon: 1e-8; beta1: 0.9; beta2: 0.99; initial learning rate: 1e-3, **Table 1**) was used to control weight updates. The loss was scaled by a factor of 1e4 prior to computing the gradients for numerical stability. To prevent potential pathological training, the GRU cell hidden states were clipped at 5 and the global gradient norm was clipped at 300.

**Table 1.** Hyperparameter ranges for RADICaL and AutoLFADS runs. Cells with a dash indicate “not applicable” for the method. LR is the learning rate. CD is the coordinated dropout rate (i.e., proportion of samples dropped at input). DO is the dropout probability for the RNN network. KL indicates the weight applied to the KL divergence of a posterior from its prior. CO indicates the controller output distributions and IC indicates the initial condition distributions. L2 indicates the weight applied to the Frobenius norm of the recurrent kernel of the GRU cell. Con indicates the controller GRU cell, Gen indicates the generator GRU cell, IC Enc indicates the initial condition encoder GRU cells. Sig Prior indicates the prior of the scaling factors applied to the sigmoid nonlinearity when mapping from factors to ZIG parameters.

|  |  |  | LR | CD | DO | KL CO | KL IC | L2 Con | L2 Gen | Sig Prior |
| --- | --- | --- | --- | --- | --- | --- | --- | --- | --- | --- |
| Defaults | RADICaL | Ranges | (1e-5, 5e-3) | (0.01, 0.99) | (0.3, 1.0) | (1e-6, 1e-4) | (1e-6, 1e-4) | (1e-5, 0.1) | (1e-5, 0.1) | (1.0, 100.0) |
|  |  | Initial values | 1e-3 | 0.5 | uniform | loguniform | loguniform | loguniform | loguniform | 20.0 |
|  |  | Explore weight | 0.3 | 0.3 | 0.3 | 0.8 | 0.8 | 0.8 | 0.8 | 0.2 |
|  |  | Limit explore | TRUE | TRUE | TRUE | FALSE | FALSE | FALSE | FALSE | FALSE |
|  | AutoLFADS | Ranges | (1e-5, 5e-3) | (0.01, 0.99) | (0.3, 1.0) | (1e-6, 1e-4) | (1e-6, 1e-4) | (1e-5, 0.1) | (1e-5, 0.1) | - |
|  |  | Initial values | 1e-3 | 0.5 | uniform | loguniform | loguniform | loguniform | loguniform | - |
|  |  | Explore weight | 0.3 | 0.3 | 0.3 | 0.8 | 0.8 | 0.8 | 0.8 | - |
|  |  | Limit explore | TRUE | TRUE | TRUE | FALSE | FALSE | FALSE | FALSE | - |

AutoLFADS is a recent implementation of the population based training (PBT) approach^58^ on LFADS to perform automatic, large-scale hyperparameter (HP) search. A detailed overview of AutoLFADS is in refs. ^17,21^. Briefly, PBT distributes training across dozens of models in parallel, and uses evolutionary algorithms to tune HPs over many generations. To do so, trials were first split into training and validation sets. At the beginning of training, the value of the searchable HPs was randomly drawn from an initial range for each individual model. At the end of each generation, a selection process was performed to choose models with higher performance (i.e., lower negative log likelihood, or NLL) on the validation set and replace the poor models with the higher performing models. The HPs of the higher performing models were perturbed before the next generation to increase the HP search space.

Training and hyperparameter search varies in the number of generations needed to converge (typically 70 - 150 generations), depending on the data and hardware used (number and type of GPUs). With our data and hardware (10x NVIDIA GeForce RTX 2080 Ti GPUs), a run of RADICaL typically converges in 3 - 5 hours. RADICaL was built in Python 2 and TensorFlow 1.14, and cloud implementations of RADICaL on Google Cloud Platform and NeuroCAAS are also being made available. Links to code and tutorials are given in *Code availability* above.

For the PBT approach, 20 single models were trained in parallel for both AutoLFADS and RADICaL runs and for both simulation and real 2P data. Generations consisted of 50 epochs, and KL and L2 regularization penalties were linearly ramped for the first 80 epochs of training during the first generation. Training was stopped when there was no improvement in performance after 25 generations. The HPs optimized by PBT were the model’s learning rate and six regularization HPs: scaling weights for the L2 penalties on the generator, controller, and initial condition encoder RNNs, scaling weights for the KL penalties on the initial conditions and controller outputs, and two dropout probabilities (“keep ratio” for coordinated dropout^21^; and RNN network dropout probability). Coordinated dropout is a regularization technique which prevents pathological overfitting by forcing the network to model only structure that is shared across neurons. The HP search ranges are detailed in **Table 1**. The magnitudes of the HP perturbation were controlled by weights and specified for different HPs (a weight of 0.3 results in perturbation factors between 0.7 and 1.3; **Table 1**). The learning rate and dropout probabilities were restricted to their specified search ranges and were sampled from uniform distributions. The KL and L2 HPs were sampled from log-uniform distributions and could be perturbed outside of the initial search ranges. Identical hyperparameter settings were used for both RADICaL and AutoLFADS and for both synthetic datasets and real 2P datasets.

RADICaL is an adaptation of AutoLFADS for 2P calcium imaging. RADICaL operates on sequences of deconvolved calcium events x(t). x(t) are modeled as a noisy observation of an underlying time-varying Zero-Inflated Gamma (ZIG) distribution^22^:

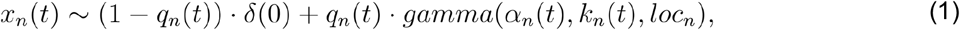

where x_n_(t) is the distribution of observed deconvolved events, a_n_(t), k_n_(t), and loc_n_ are the scale, shape, and location parameters, respectively, of the gamma distribution, and q_n_(t) denotes the probability of non-zeros, for neuron *n* at time *t*. loc_n_ was fixed as the minimum nonzero deconvolved event (s_min_). In the original AutoLFADS model, factors were mapped to a single time-varying parameter for each neuron (the Poisson firing rate) via a linear transformation followed by an exponential nonlinearity. RADICaL instead infers the three time-varying parameters for each neuron, a_n_(t), k_n_(t), and q_n_(t), by linearly transforming the factors followed by a trainable scaled sigmoid nonlinearity (sig_n_). sig_n_ is a positive parameter that scales the outputs of the sigmoid to be in a range between 0 and sig_n_, and is optimized alongside network weights. An L2 penalty is applied between sig_n_ and a PBT-searchable prior (**Table 1**) to prevent extreme values. The training objective is to minimize the negative log-likelihood of the deconvolved events given the inferred parameters:

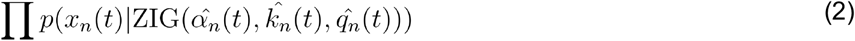

The event rate for neuron *n* at time *t* was estimated by taking the mean of the inferred ZIG distribution:

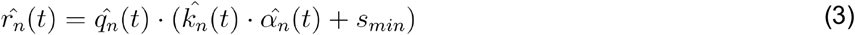

RADICaL uses an SBTT training strategy to achieve sub-frame modeling resolution. RADICaL operates on binned deconvolved calcium events, with bin size smaller than the frame timebase of imaging. Bins where the neurons were sampled were filled with the corresponding event rates, while bins where the neurons were not sampled were filled with NaNs. Choosing the sub-frame bin width involves a trade-off. Finer bins improve the possible temporal resolution, but if the data are binned too finely, there may be very few neurons in certain bins, leading to uncertainty about the estimated latent states. It is important to choose the sub-frame bin size to ensure a reasonable number of neurons in each bin. We recommend a neuron count greater than 20 per sub-frame bin based on the results from our neuron downsampling experiments.

The networks output the time-varying ZIG distribution at each sub-frame timestep; however, a mask was applied to the timesteps where the NaN samples were to prevent the cost computed from these timesteps being backpropagated during gradient calculation. As a result, the model weights were only updated based on the cost at the sampled timesteps. The reconstruction cost also excluded the cost calculated at the non-sampled timesteps so the PBT model selection was not affected by the cost computed from the non-sampled timesteps.

### Simulation experiments

#### Generating spike trains from an underlying Lorenz system

Synthetic data were generated using the Lorenz system as described in the original LFADS work^15,16^. Lorenz parameters were set to standard values (σ: 10, ρ: 28, and β: 8/3), and Δt was set to 0.01. Datasets with different speeds of dynamics were generated by downsampling the original generated Lorenz states by different factors. The speed of the Lorenz dynamics was quantified based on the peak location of the power spectra of the Lorenz Z dimension, with a sampling frequency of 100 Hz. The downsampling factors were 3, 5, 7, 9, 11 and 14 for speeds 4, 7, 10, 13, 15 and 20 Hz, respectively. Each dataset/speed consisted of 8 conditions, with 60 trials per condition. Each condition was obtained by starting the Lorenz system with a random initial state vector and running it for 900 ms. The trial length for the 4 Hz dataset was longer (1200 ms) than that of other datasets (900 ms) to ensure that all conditions had significant features to be modeled - with shorter windows, the extremely low frequency oscillations caused the Lorenz states for some conditions to have little variance across the entire window, making it trivial to approximate the essentially flat firing rates. We simulated a population of 278 neurons with firing rates given by linear readouts of the Lorenz state variables using random weights, followed by an exponential nonlinearity. Scaling factors were applied so the baseline firing rate for all neurons was 3 spikes/sec. Each bin represents 10 ms and an arbitrary frame time was set to be 30 ms (i.e., one “imaging frame” takes 3 bins). Spikes from the firing rates were then generated by a Poisson process.

#### Generating fluorescence signals from synthetic spike trains

Realistic fluorescence signals were generated from the spike trains by convolving them with a kernel for an autoregressive process of order 2 and passing the results through a nonlinearity that matched values extracted from the literature for the calcium indicator GCaMP6f^5,59^ (**Supp Fig. 2a & b**). Three noise sources were added to reproduce variability present in real data^60–62^: Gaussian noise to the size of the calcium spike, and Gaussian and Poisson noise to the final trace (**Supp Fig. 2a & b**). This fluorescence generation process was realized as follows: First, spike trains s(t) were generated from the Lorenz system as mentioned above. Independent Gaussian noise (sd = 0.1) was added to each spike in the spike train to model the variability in spike amplitude. Next, we modeled the calcium concentration dynamics c(t) as an autoregressive process of order 2:

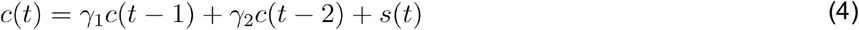

with s(t) representing the number of spikes at time t. The autoregressive coefficients *γ*_1_ and *γ*_2_ were computed based on the rise time, decay time (*τ_on_* = 20 ms, *τ_off_* = 400 ms for GCaMP6f) of the calcium indicators, and the sampling frequency. Note that while there is substantial variability in taus across neurons in real data^5^, selecting and mimicking this variability was not relevant in our work, because we compared the methods (i.e., RADICaL, AutoLFADS, and s-deconv) after deconvolution. The calcium concentration dynamics were further normalized so that the peak height of the calcium dynamics generated from a single spike equalled one, regardless of the sampling frequency. Subsequently, we computed the noiseless fluorescence signals by passing the calcium dynamics through a nonlinear transformation estimated from the literature^59^ for the calcium indicator GCaMP6f (**Supp Fig. 2c & d**). After the nonlinear transformation, the relationship between spike size and trace size was corrupted, and therefore we assumed the baseline of fluorescence signals to be zero and the signals were rescaled to the range in [0,1] using min-max normalization. Finally, Gaussian noise (~N(0,sn)) and Poisson noise (simulated as gaussian with mean 0 and variance proportional to the signal amplitude at each time point via a constant *d*) were added to the normalized traces. The resulting fluorescence traces had the same sampling frequency as the synthetic spike trains (100 Hz).

A crucial parameter is the noise level associated with each fluorescence trace. High noise levels lead to very poor spike detection and very low noise levels enable a near-perfect reconstruction of the spike train. In order to select a realistic level of noise we matched the correlations between real and inferred spike trains of the simulated data to those observed in a recent benchmarking study^13^. We found that a truncated normal distribution of noise level for Gaussian and Poisson noise best matched the correlations. More specifically, for each neuron, *sn*=*d* was sampled independently from a truncated normal distribution N(0.12, 0.02) with the tail below 0.06 removed. With the above noise setting, the mean correlation coefficient *r* between the deconvolved events and ground truth spikes was 0.32, which is consistent with the standard results reported in the “spikefinder” paper^13^ for OASIS. In our additional tests of model tolerance to spike inference noise, the Gaussian noise added to the fluorescence traces was increased by 2x or 4x. It is worth stressing that real data feature a broad range of noise levels that depend on the imaging conditions, depth, expression level, laser power and other factors. Here we did not attempt to investigate all possible noise conditions, but instead, we aimed to create a simulation with known latent variables (i.e., low-dimensional factors and event rates) that reasonably approximated realistic signal-to-noise levels, in order to provide a tractable test case to compare RADICaL to other methods before attempting comparisons on real data.

#### Recreating variability in sampling times due to 2p laser scanning

The fluorescence traces were simulated at 100 Hz as mentioned above. A subsampling step was then performed with sampling times for each neuron staggered in time to simulate the variability in sampling times due to 2p laser scanning (as in **Fig. 1e**). This produced fluorescence traces where individual neurons were sampled at 33.3 Hz, with phases of 0, 11, 22 ms based on each neuron’s location (top, middle and bottom of the FOV, respectively). To break this down, each neuron was sparsely sampled every three time points and the relative sampled times between neurons were fixed. For example, in trial 1, neuron 1 was sampled at time points 1, 4, 7, … and neuron 2 was sampled at time points 2, 5, 8, …; in trial 2, neuron 1 was sampled at time points 2, 5, 8, … and neuron 2 was sampled at time points 3, 6, 9, …. Thus, the sampling frequency for each individual neuron was 33.3 Hz, while the sampling frequency for the population was retained at 100 Hz by filling the non-sampled time points with NaNs. The resulting 33.3 Hz simulated fluorescence signals for each individual neuron (i.e., with NaNs excluded) were deconvolved using OASIS^25^ (as implemented in CaImAn^26^) using an auto-regressive model of order 1 with s_min_ of 0.1. For experiments with slower imaging speeds, the same steps were repeated but the simulated 100 Hz fluorescence signals were subsampled at different rates (i.e., 16 Hz, 8 Hz and 2 Hz).

#### Data preparation for each method

Four methods (RADICaL, AutoLFADS, s-deconv and s-sim-fluor) were compared by their performance on recovering the ground truth latent states across different datasets/speeds. Trials (480 total for each simulated dataset) were split into 80/20 training and validation sets for modeling AutoLFADS and RADICaL. To prepare data for non-RADICaL methods, non-sampled bins were removed so all the sampled bins were treated as if they were sampled at the same time and each bin then represented 30 ms (i.e., sampling frequency = 33.3 Hz). Preparing the data for AutoLFADS required discretizing the deconvolved events into spike count estimates, because AutoLFADS was primarily designed to model discrete spiking data. In the discretizing step, if the event rate was 0, it was left as 0; if the event rate was between 0 and 2, it was cast to 1 (to bias toward the generally higher probability of fewer spikes). If the event rate was greater than 2, it was rounded down to the nearest integer. We note that this is one of many possible patches to convert continuously-valued event intensities to natural numbers for compatibility with the Poisson distribution and AutoLFADS; a more principled solution would be to modify the network to use the ZIG distribution, as we have done in RADICaL. With s-deconv, the deconvolved events were smoothed by convolution with a Gaussian filter (6 ms s.d.) to produce event rates. With s-sim-fluor, the generated fluorescence signals were smoothed by convolution with a Gaussian filter (6 ms s.d.) to produce event rates. The choice of filter width was optimized by sweeping values ranging from 3 to 40 ms. Smoothing with a 6 ms s.d. filter gave the highest performance in recovering the ground truth Lorenz states for experiments with higher Lorenz frequencies (i.e., >= 10 Hz). The event rates produced from RADICaL had a sampling frequency of 100 Hz, while the event rates produced from the non-RADICaL methods had a sampling frequency of 33.3 Hz. The non-RADICaL rates were then resampled at 100 Hz using linear interpolation.

#### Mapping to ground truth Lorenz states

Since our goal was to quantify modeling performance by estimating the underlying Lorenz states, we trained a mapping from the output of each model (i.e., the event rates) to the ground truth Lorenz states using ridge regression. First, we split the trials into training (80%) and test (20%) sets. We used the training set to optimize the regularization coefficient using 5-fold cross-validation, and used the optimal regularization coefficient to train the mapping on the full training set. We then quantified state estimation performance by applying this trained mapping to the test set and calculating the coefficient of determination (*R^2^*) between the true and predicted Lorenz states. We repeated the above procedure five times with train/test splits drawn from the data in a complementary fashion. We reported the mean *R^2^* across the repeats, such that all reported numbers reflect held-out performance. We tested whether the difference of *R^2^* between each pair of methods was significant by performing a paired, one-sided Student’s t-test on the distribution of *R^2^* across the five folds of predictions. In our simulations we observed a delay caused by deconvolution, where the deconvolved events came systematically later than the true spikes, consistent with findings in a recent study^41^. We swept across different lags between the event rates and the true latent states in the latent mapping analysis and empirically found that correcting for a delay of 30 ms gave the highest latent recovery performance across the methods (data not shown). We therefore included a 30 ms lag correction in all of our latent mapping analyses.

#### Additional tests of deconvolution using MLspike

To test whether RADICaL works on deconvolved events that have a spike-time-like structure, we tested MLspike^27^ as an alternative for deconvolution. Calcium traces were generated using the identical steps as described above. For MLspike, the cubic polynomial model was chosen as the nonlinearity model consistent with GCaMP6f. The drift parameter was set to 0.001. The decay time constant tau was set to 0.4s. We did not use auto calibration in MLspike because it produced inconsistent results in our tests. Instead, to give MLspike the best chance at high performance, we manually tuned the remaining parameters in MLspike by reducing the error rates for inferred spikes compared to ground truth spikes using a small subset of neurons. Transient amplitude was set to 1 and the noise parameter sigma was set to 0.15. Spikes inferred by MLspike were then prepared for AutoLFADS and RADICaL as described above. Note that the discretizing step was omitted here when preparing data for AutoLFADS.

### Real 2p experiments

#### Surgical procedures

All procedures were approved by the University of Chicago Animal Care and Use Committee. Two male Ai148D transgenic mice (TIT2L-GC6f-ICL-tTA2, stock 030328; Jackson Laboratory) were used. Each mouse underwent a single surgery. Mice were injected subcutaneously with dexamethasone (8 mg/kg) 24 hours and 1 hour before surgery. Mice were anesthetized with 2-2.5% inhaled isoflurane gas, then injected intraperitoneally with a ketamine-medetomidine solution (60 mg/kg ketamine, 0.25 mg/kg medetomidine), and maintained on a low level of supplemental isoflurane (0-1%) if they showed any signs that the depth of anesthesia was insufficient. Meloxicam was also administered subcutaneously (2 mg/kg) at the beginning of the surgery and for 1-3 subsequent days. The scalp was shaved, cleaned, and resected, the skull was cleaned and the wound margins glued to the skull with tissue glue (VetBond, 3M), and a 3 mm circular craniotomy was made with a 3 mm biopsy punch centered over the left CFA/S1 border. The coordinates for the center of CFA were taken to be 0.4 mm anterior and 1.6 mm lateral of bregma. The craniotomy was cleaned with SurgiFoam (Ethicon) soaked in phosphate-buffered solution (PBS), then virus (AAV9-CaMKII-Cre, stock 2.1*10^13^ particles/nL, 1:1 dilution in PBS, Addgene) was pressure injected (NanoJect III, Drummond Scientific) at two or four sites near the target site, with 140 nL injected at each of two depths per site (250 and 500 μm below the pia) over 5 minutes each. The craniotomy was then sealed with a custom cylindrical glass plug (3 mm diameter, 660 μm depth; Tower Optical) bonded (Norland Optical Adhesive 61, Norland) to a 4 mm #1 round coverslip (Harvard Apparatus), glued in place first with tissue glue (VetBond) and then with cyanoacrylate glue (Krazy Glue) mixed with dental acrylic powder (Ortho Jet; Lang Dental). A small craniotomy was also made using a dental drill over right CFA at 0.4 mm anterior and 1.6 mm lateral of bregma, where 140 nL of AAVretro-tdTomato (stock 1.02*10^13^ particles/nL, Addgene) was injected at 300 μm below the pia. This injection labeled cells in left CFA projecting to the contralateral CFA. Here, this labeling was used solely for stabilizing the imaging plane (see below). The small craniotomy was sealed with a drop of Kwik-Cast (World Precision Instruments). Two layers of MetaBond (C & B) were applied, then a custom laser-cut titanium head bar was affixed to the skull with black dental acrylic. Animals were awoken by administering atipamezole via intraperitoneal injection and allowed to recover at least 3 days before water restriction.

#### Behavioral task

The behavioral task (Fig. 3a) was a variant of the water reaching task of ref. ^28^ which we term the “water grab” task. This task was performed by water-restricted, head-fixed mice, with the forepaws beginning on paw rests (eyelet screws) and the hindpaws and body supported by a custom 3D printed clear acrylic tube enclosure. After holding the paw rests for 700-900 ms, a tone was played by stereo speakers and a 2-3 μL droplet of water appeared at one of two water spouts (22 gauge, 90-degree bent, 1” blunt dispensing needles, McMaster) positioned on either side of the snout. The pitch of the tone indicated the location of the water, with a 4000 Hz tone indicating left and a 7000 Hz tone indicating right, and it lasted 500 ms or until the mouse made contact with the correct water spout. The mouse could grab the water droplet and bring it to its mouth to drink any time after the tone began. Both the paw rests and spouts were wired with capacitive touch sensors (Teensy 3.2, PJRC). Good contact with the correct spout produced an inter-trial interval of 3-6 s, while failure to make contact (or insufficiently strong contact) with the spout produced an inter-trial interval of 20 s. Because the touch sensors required good contact from the paw, this setup encouraged complex contacts with the spouts. The mice were trained to make all reaches with the right paw and to keep the left paw on the paw rest during reaching. Training took approximately two weeks, though the behavior continued to solidify for at least two more weeks. Data presented here were collected after 6-8 weeks’ experience with the task. Control software was custom written in MATLAB R2018a using PsychToolbox 3.0.14, and for the Teensy. Touch event monitoring and task control were performed at 60 Hz.

Behavior was also recorded using a pair of cameras (BFS-U3-16S2M-CS, FLIR; varifocal lenses COZ2813CSIR2, Computar) mounted 150 mm from the right paw rest at 10° apart to enable 3D triangulation. Infrared illuminators enabled behavioral imaging while performing 2p imaging in a darkened microscope enclosure. Cameras were synchronized and recorded at 150 frames per second with real-time image cropping and JPEG compression, and streamed to one HDF5 file per camera (areaDetector module of EPICS, CARS). The knuckles and wrist of the reaching paw were tracked in each camera using DeepLabCut^29^ and triangulated into 3D using camera calibration parameters obtained from the MATLAB Stereo Camera Calibration toolbox^63,64^. To screen the tracked markers for quality we created distributions of all inter-marker distances in 3D across every labeled frame and identified as problematic frames with any inter-marker distance exceeding the 99.9th percentile of its respective distribution. Trials with more than one problematic frame in the period of −200 ms to 800 ms after the raw reach onset were discarded (where reach onset was taken as the first 60 Hz tick after the paw rest touch sensor fell below contact threshold). The kinematics of all trials that passed this screening procedure were visualized to confirm quality. Centroid marker kinematics were obtained by averaging the kinematics of all paw markers, locking them to behavioral events and then smoothing using a Gaussian filter (15 ms s.d.). To obtain velocity and acceleration, centroid data was numerically differentiated with MATLAB’s *diff* function and then smoothed again using a Gaussian filter (15 ms s.d.).

#### Two-photon imaging

Calcium imaging was performed with a Neurolabware two-photon microscope and pulsed Ti:sapphire laser (Vision II, Coherent). Depth stability of the imaging plane was maintained using a custom plugin that acquired an image stack at the beginning of the session (1.4 μm spacing), then compared a registered rolling average of the red-channel data to each plane of the stack. If sufficient evidence indicated that a plane not at the center of the stack was a better match to the image being acquired, the objective was automatically moved to compensate. This typically resulted in a slow and steady upward (outward) movement of the objective over the course of the session. This plane drift is probably due to ETL warming, as it occurred when imaging slides at high power but not low power. The power range used in imaging was approximately 50-65 mW average power, including the net power reduction due to end-of-line blanking.

Offline, images were run through Suite2p to perform motion correction, region-of-interest (ROI) detection, and fluorescence extraction from both ROIs and neuropil. ROIs were manually curated using the Suite2p GUI to retain only those corresponding to somas. We then subtracted the neuropil signal scaled by 0.7^7^. Neuropil-subtracted ROI fluorescence was then detrended by performing a running 10th percentile operation, smoothing with a Gaussian filter (20 s s.d.), then subtracting the result from the trace. This result was fed into OASIS^25^ using the ‘thresholded’ method, AR1 event model, and limiting the tau parameter to be between 300 and 800 ms. Neurons were discarded if they did not meet a minimum signal-to-noise (SNR) criterion. To compute SNR, we took the fluorescence at each time point when OASIS identified an “event” (non-zero), computed (fluorescence - neuropil) / neuropil, and computed the median of the resulting distribution. ROIs were excluded if this value was less than 0.05. To put events on a more useful scaling, for each ROI we found the distribution of event sizes, smoothed the distribution (ksdensity in MATLAB, with an Epanechnikov kernel and log transform), found the peak of the smoothed distribution, and divided all event sizes by this value. This rescales the peak of the distribution to have a value of unity. Data from two mice and two brain areas (4 sessions in total) were used (Mouse1/M1: 510 neurons, 560 trials; Mouse1/S1: 543 neurons, 506 trials; Mouse2/M1: 439 neurons, 475 trials; Mouse2/S1: 509 neurons, 421 trials).

#### Data preparation for modeling with RADICaL and AutoLFADS

To prepare data for RADICaL, the deconvolved events were normalized by the s_min value output by OASIS so that the minimal event size was 0.1 across all neurons. The deconvolved events for individual neurons had a sampling rate equal to the frame rate (31.08 Hz). For modeling with RADICaL, the deconvolved events were assigned into 10ms bins using the timing of individual measurements for each neuron to achieve sub-frame resolution (i.e., 100 Hz). The non-sampled bins were filled with NaNs. To prepare data for AutoLFADS, the deconvolved events were rescaled using the distribution-scaling method described above, and casted using the casting step described in the simulation section. For both AutoLFADS and s-deconv, the deconvolved events were assigned into a single time bin per frame (i.e., 32.17 ms bins) to mimic standard processing of 2p imaging data, where the sub-frame timing of individual measurements is discarded. Trials were created by aligning the data to 200 ms before and 800 ms after reach onset (100 time points per trial for RADICaL, and 31 time points per trial for AutoLFADS and s-deconv). An individual RADICaL model and AutoLFADS model were trained for each dataset (4 total). Failed trials (latency to contact with correct spout > 15 s for Mouse1, 20 s for Mouse2), or trials where the grab to the incorrect spout occurred before the grab to the correct spout, were discarded. For each dataset, trials (Mouse1/M1: 552 total; Mouse1/S1: 500 total; Mouse2/M1: 467 total; Mouse2/S1: 413 total) were split into 80/20 training and validation.

#### Trial grouping

PSTH analysis and low dimensional neural trajectory visualization were performed based on subgroups of trials. Trials were sorted into two subgroups per spout based on the Z dimension (height) of hand position. The hand position was obtained by smoothing the centroid marker position with a Gaussian filter (40 ms s.d.). Time windows where the height of hand was used to split trials were hand-selected to present a good separation between subgroups of hand trajectories. For Mouse1/M1, a window of 30 ms to 50 ms after reach onset was used to split left condition trials and a window of 180 ms to 200 ms after reach onset was used to split right condition trials; for Mouse1/S1, a window of 140 ms to 160 ms after reach onset was used to split both left and right condition trials; for both Mouse2/M1 and Mouse2/S1, a window of 30 ms to 50 ms after reach onset was used to split both left and right condition trials. For both left or right conditions and for all mice/areas, 55 trials with the lowest and highest heights were selected as group 1 and group 2, respectively; trials with middle-range heights were discarded.

#### PSTH analysis and comparing RADICaL and AutoLFADS single-trial rates

RADICaL was first validated by comparing the PSTHs computed using RADICaL inferred event rates and the empirical PSTHs. Empirical PSTHs were computed by trial-averaging s-deconv rates (40 ms kernel s.d., 32.17 ms bins) within each of the 4 subgroups of trials. RADICaL inferred rates were first downsampled from 100 Hz to 31.08 Hz with an antialiasing filter applied, to match the sampling frequency (i.e., the frame rate) of the original deconvolved signals. RADICaL PSTHs were computed by similarly averaging RADICaL rates. Single-trial inferred rates were then compared to the empirical PSTHs to assess how well each method recapitulated the empirical PSTHs on single trials. The correlation coefficient (r) was computed between inferred single-trial event rates and the corresponding empirical PSTHs in a cross-validated fashion, i.e., each trial’s inferred event rate was compared against an empirical PSTH computed using all other trials within the subgroup. *r* was assessed for the time window spanning 200 ms before to 800 ms after reach onset, and computed by concatenating all trials across the four subgroups, yielding one *r* for each neuron. Neurons that had fewer than 40 nonzero events within this time window (across all trials) were excluded from the analysis.

#### Low-D analysis

To visualize the low-dimensional neural trajectories that RADICaL produced, principal component analysis (PCA) was performed on RADICaL inferred rates. RADICaL rates (aligned to 200 ms before and 800 ms after reach onset) were log-transformed (with 1e-4 added to prevent numerical precision issues) and normalized to have zero mean and unit standard deviation for each neuron. PCA was applied to the trial-averaged rates and the projection matrix was then used to project the log-transformed and normalized single-trial rates (aligned to 200 ms before and 400 ms after reach onset) onto the top 3 PCs.

#### Subgroup distance ratio analysis

To quantitatively measure how informative RADICaL was about the subgroup identity of each trial, a subgroup distance ratio analysis was performed in the inferred rate space. For each trial at each time point, we measured the Euclidean distances to the corresponding time point of each other trials within the same subgroup as well as the distances to the corresponding time point of each trial from the other subgroup of the same condition. The distance ratio was computed as the ratio of the mean across-subgroup differences to the mean within-subgroup distances. A distance ratio greater than one indicates that the trial is more closely grouped with the trials within the same subgroup compared to the other subgroup. An averaged distance ratio was computed across all trials for each time point.

#### Decoding analysis

RADICaL-inferred rates, AutoLFADS-inferred rates, and s-deconv (Gaussian kernel 40 ms s.d.) rates were used to decode hand position and velocity using ridge regression. The hand position and velocity were obtained as described above and binned at 10 ms (i.e., 100 Hz). The non-RADICaL rates were retained to a sampling frequency of 100 Hz using linear interpolation. For simplicity, we did not include a lag between the neural data and kinematics. However, additional analyses confirmed that adding a lag did not alter the results (data not shown). Trials with an interval between water presentation and reach onset that was longer than a threshold were discarded due to potential variations in behavior (e.g., inattention). The threshold was selected arbitrarily for different sessions based on the actual distribution of the intervals in the session (Mouse1/M1: 500 ms; Mouse1/S1: 600 ms; Mouse2/M1: 400 ms; Mouse2/S1: 600 ms). The data were aligned to 50 ms before and 350 ms after reach onset. The decoder was trained and tested using cross-validated ridge regression. First, we split the trials into training (80%) and test (20%) sets. We used the training set to optimize the regularization coefficient using 5-fold cross-validation, and used the optimal regularization coefficient to train the decoder on the full training set. This trained decoder was applied to the test set, and the coefficient of determination (*R^2^*) was computed and averaged across x-, y- and z-kinematics. We repeated the above procedure five times with train/test splits drawn from the data in an interleaved fashion. We reported the mean *R^2^* across the repeats, such that all reported numbers reflect held-out performance. We tested whether the difference of *R^2^* between each pair of methods was significant by performing paired, one-sided Student’s t-Tests on the distribution of *R^2^* across the five folds of predictions.

One possible concern is that RADICaL improves decoding not because the single-trial traces are better denoised, but instead because they for some reason result in learning a better decoder. To address this, we performed a “cross-decoder” analysis where the decoder trained with s-deconv rates was applied to the RADICaL inferred rates. Note that it is not guaranteed that the cross-decoder would give better performance even if RADICaL’s rates are better denoised, because this is also a task of generalization - during training, the decoder did not see the RADICaL rates which might have different distributions of signal-to-noise across neurons or might require a different level of regularization. Despite this being a difficult task, the cross-decoder analysis shows improved performance over the original s-deconv decoding (**Supp. Fig. 21**). This suggests that the improvement seen in **Fig. 4a & b** does not merely reflect the training performance of the decoder but also demonstrates the higher quality of the inferred rates themselves.

#### Coherence analysis

Coherence was computed between the true and predicted kinematics (window: 200 ms before and 500 ms after reach onset) across all trials and across all x-, y- and z-dimensions using magnitude-squared coherence (MATLAB: mscohere). The power spectral density estimation parameters within mscohere were specified to ensure a robust calculation on the single trial activity: Hanning windows with 35 timesteps (i.e., 350 ms) for the FFT and window size, and 25 timesteps (i.e., 250 ms) of overlap between windows.

Although the coherence analysis presents the performance of each method as a function of frequency (**Fig. 4c**), the values are not directly comparable to the latent recovery analysis in simulation (**Fig. 2c**). In the simulations, the known, true underlying latent states can be used to directly measure success. In contrast, with real data the true underlying latent states are unknown and the behavioral measurements (hand position and velocity) are indirect correlates. The coherence metric therefore includes other sources of error such as muscle and tracking noise. Both the quicker drop as frequency increases, and the smaller difference between methods, could potentially be explained by the limitations of indirect measurement. In addition, the relationship between neural activity and hand position/velocity may be nonlinear or history-dependent, while our decoding was linear and instantaneous.

#### Reaction time prediction analysis

RADICaL-inferred rates, AutoLFADS-inferred rates, and s-deconv (Gaussian kernel 40 ms s.d.) rates were used to predict reaction time (RT) using logistic regression. This analysis follows the same procedure used in ref. ^30^. Reaction time was defined as the interval from water presentation to movement onset. Movement onset was defined as the time when the speed of the paw centroid exceeded 20% of this trial’s peak speed. Single-trial rates by the three methods were first aligned to movement onset, then projected into the top 10-PC space. Data were binned into a “premovement” time point (100ms before to movement onset) and a “movement” time point (movement onset to 100ms after). Trials were split into training (75%) and test (25%) sets. A logistic regression classifier was trained using the training set and returned a projection dimension that best discriminated between premovement and movement data. The projection returned by logistic regression was then used to project the test trials binned at original bin size (i.e., 100 Hz). The RT was predicted as the time when the projected activity crossed a 50% threshold. The correlation coefficient (*r*) was computed between the true and predicted RTs for the test trials, such that the reported numbers reflect held-out performance.

#### t-SNE analysis on the weights mapping from factors to ZIG parameters

RADICaL relies on sub-frame bins in which neurons are grouped based on their spatial locations within the FOV. Because this strategy results in consistent neuron grouping, it could potentially result in different groups of neurons corresponding to different latent factors. To test whether such an artifact existed, we visualized the transformation from latents to neurons by using t-SNE to reduce the 300-dimensional weights vector (100 factors * 3 ZIG parameters) into a 2-D t-SNE space for each individual neuron (510 neurons total) (**Supp. Fig. 22**). We did not observe a relationship between neurons’ position within the field of view (i.e., top, middle, and bottom) and the underlying factors. This suggested that the model did not use distinct factors for sets of neurons that were sampled with different phases, despite neurons in distant portions of the FOV never being grouped in the same bin.

#### Neuron downsampling

Two neuron downsampling experiments were performed with different procedures to test the methods’ tolerance to low neuron counts. The first procedure was designed to mimic scanning a sparse population of neurons. To do so, the number of neurons included when training RADICaL or AutoLFADS was gradually reduced by randomly dropping a subset of neurons from the previous subset, with a fraction kept of 1, 3/4, 1/2, 1/4, 1/8 or 1/16. This results in 439, 329, 219, 109, 54 or 27 neurons kept for the Mouse2/M1 dataset, and 543, 407, 271, 135, 67 or 33 kept for the Mouse1/S1 dataset. One RADICaL model and one AutoLFADS model were trained for each number of neurons. Decoding was performed using ridge regression (see above).

The other procedure was designed to emulate scanning a smaller field of view, such as when using a lens relay to image deep structures., Here, the number of neurons included when training RADICaL or AutoLFADS was gradually reduced by limiting the area of FOV that the neurons were sampled from. The area was shrunk from the entire FOV with an area-to-FOV ratio of 1, 25/36, 9/16, 1/4, and 1/9, resulting in the number of included neurons being 439, 321, 262, 121 or 59 for Mouse2/M1. An individual RADICaL model and AutoLFADS model were trained for each number of neurons. Decoding was performed using ridge regression (see above). Note that this analysis represents a lower bound on performance: for this proof-of-concept, we simply artificially excluded data from outside the restricted FOVs, which resulted in substantial time periods that lacked data entirely (e.g., 2/3 of the total sampling time for the smallest FOV considered). In a real application, those time periods that were artificially excluded could instead be used to monitor other brain areas or layers, or to monitor the same neurons with higher sampling rates, either of which might be expected to provide additional information.

