## Supplementary Material for "A deep learning framework for inference of single-trial neural population dynamics from calcium imaging with sub-frame temporal resolution"

**Supplementary materials:**

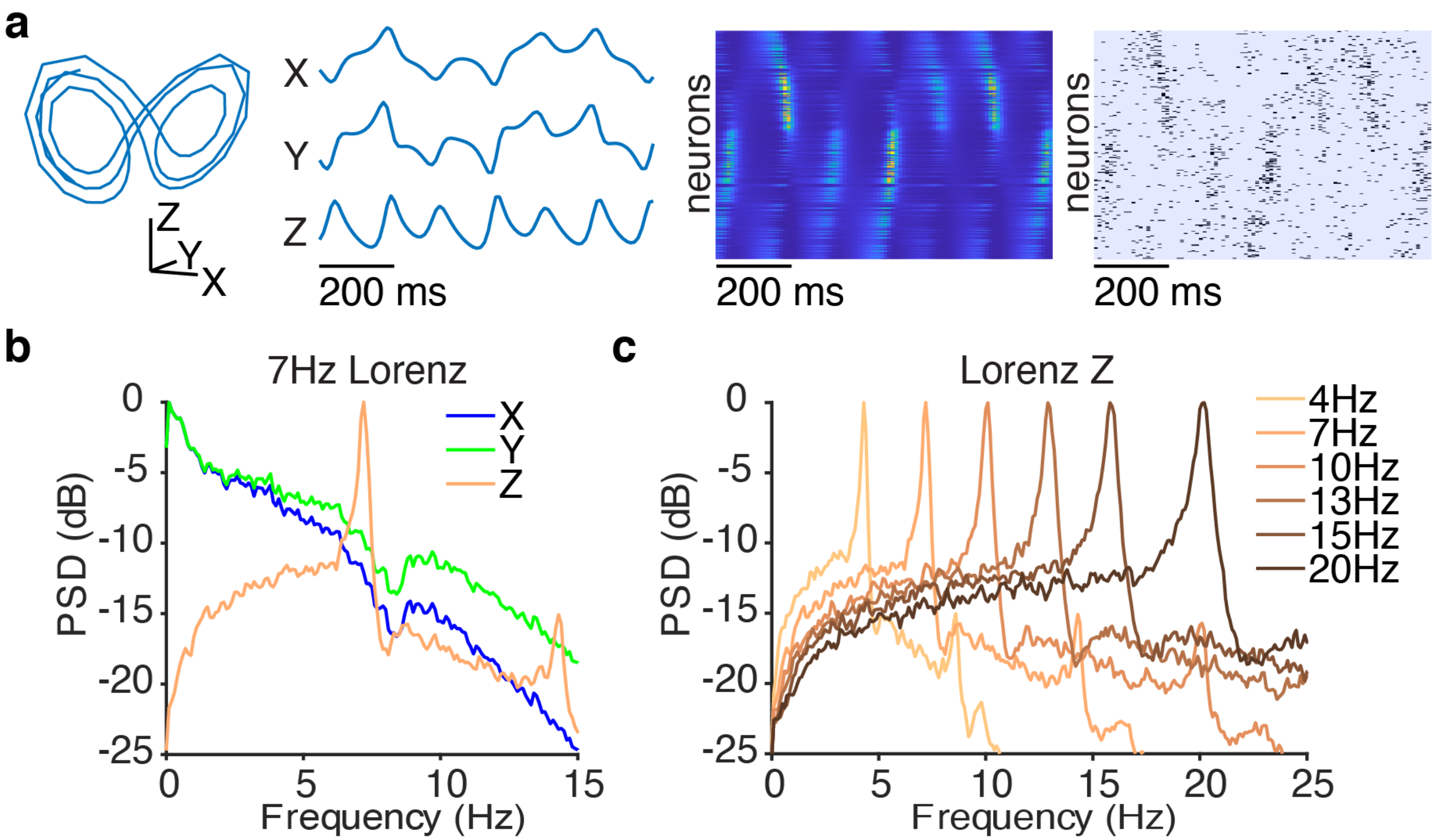


**Supplementary Figure 1 | Simulation of Lorenz system at different speeds.** This figure illustrates the underlying dynamical system used for the simulation experiments. (a) An example Lorenz trajectory in a 3-dimensional state space (far left) and with three dynamic variables plotted as a function of time (middle left) for a system with *Z*-oscillation peak frequency of 7 Hz (i.e., the power spectrum of the Lorenz system's Z-dimension had a pronounced peak at 7 Hz). Firing rates for the simulated neurons were computed by a linear readout of the Lorenz variables followed by an exponential nonlinearity (middle right). Spikes from the firing rates were then generated by a Poisson process (far right). The example trial shown here is identical to "Trial 2" in **Fig. 2a**, but with a wider plotting window. (b) Power spectrum of the individual Lorenz variables for the system with a *Z*-oscillation peak frequency at 7 Hz. Because only the Z variable has a clear peak in the power spectrum, this variable was used exclusively for all further analyses in simulations except Supplementary Figure 3. (c) Power spectrum of the *Z* dimension for Lorenz systems simulated with different *Z*-oscillation peak frequencies.

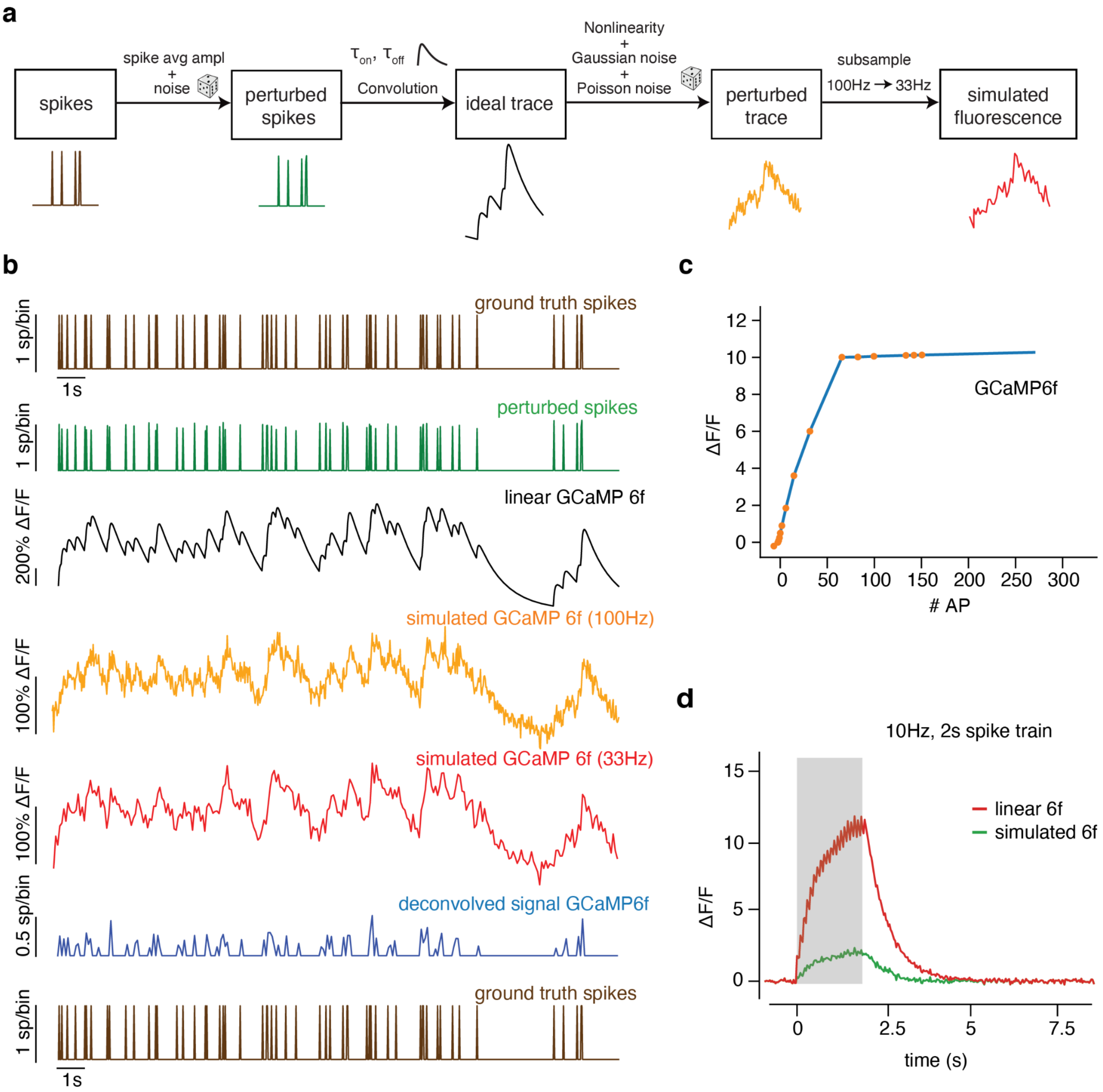


**Supplementary Figure 2 | Simulation pipeline to generate artificial fluorescence traces from the underlying Lorenz system.** (a) This pipeline begins from the Poisson-random spikes generated in the far-right panel of Supplementary Figure 1. Calcium traces were generated by first corrupting the spikes with amplitude noise, then modeling the dynamics of calcium indicators in response to a spike with an autoregressive process of order 2 transformed by a piecewise-linear non-linearity. Sources of noise corrupting this fluorescence trace were then added. The nonlinearity and noise sources were chosen to approximate the variability observed in real data. (b) Example ground truth and simulated data using a GCaMP6f model. From top to bottom: original ground truth spikes fed into the simulator, perturbed spikes, idealized calcium trace, fluorescence trace with nonlinearity and noise sources added, fluorescence trace after subsampling, deconvolved spikes, and finally original ground truth spikes fed into the simulator (shown again for comparison; same as top). (c) Estimated nonlinearities for GCaMP6f from ref. [59]. (d) Example traces generated by the simulator for a train of 10 Hz stimuli, with and without nonlinearity applied.

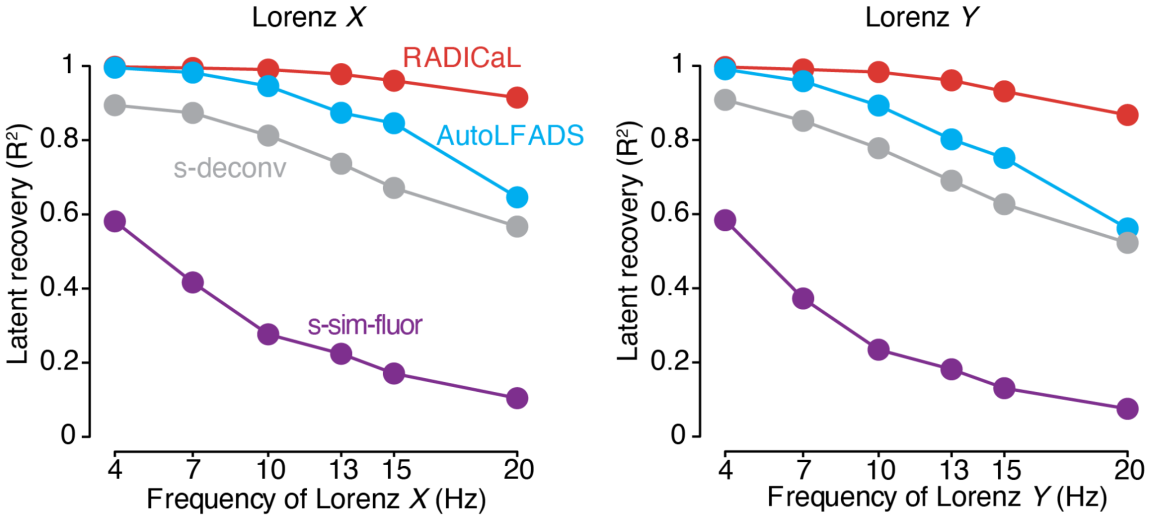


**Supplementary Figure 3 | Performance of estimating other Lorenz dimensions in the simulation experiments.** Performance of estimating Lorenz *X* (*left*) and *Y* (*right*) dimensions as a function of simulation frequency was quantified by variance explained ($R^2$) for all 4 methods. Note that these variables are dominated by lower frequencies than the *Z* variable used in other figures, and therefore make for an easier challenge. We therefore used the *Z* variable for all other results.

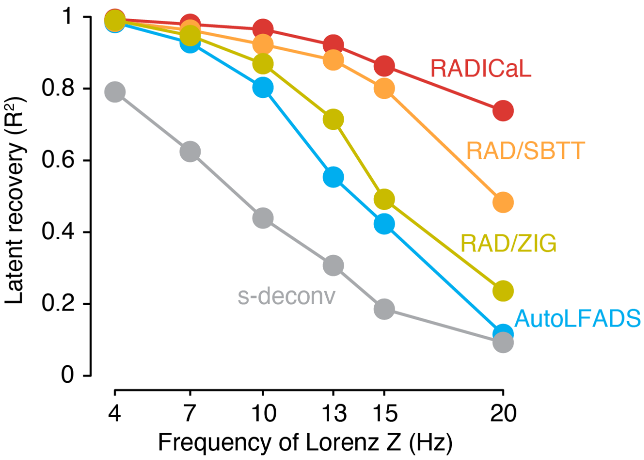


**Supplementary Figure 4 | Both SBTT and ZIG improve latent recovery performance separately.** To understand the contributions of ZIG and SBTT independently in RADICaL's performance in latent recovery, we fit RADICaL to different Lorenz oscillation frequencies with only the ZIG emission model enabled (no SBTT; "RAD/ZIG") or only SBTT enabled (no ZIG; "RAD/SBTT"). Performance in estimating the Lorenz *Z* dimension as a function of Lorenz oscillation frequency was quantified by variance explained ($R^2$). RAD/ZIG performed a little better than AutoLFADs, while RAD/SBTT performs substantially better, but combining both (the full RADICaL model) performed substantially better still.

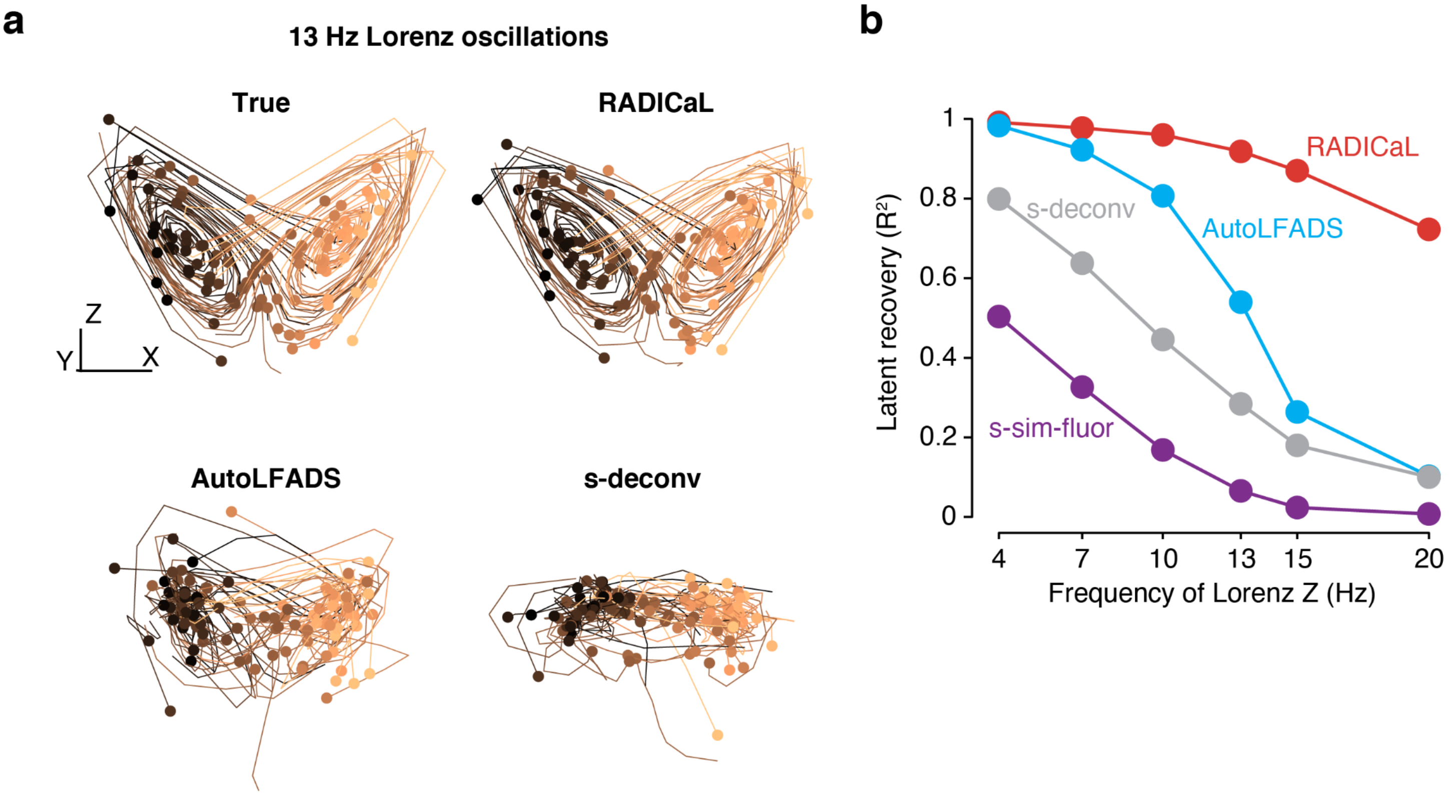


**Supplementary Figure 5 | RADICaL retains high latent recovery performance in a simulation experiment that lacks stereotyped conditions.** This analysis was targeted at determining whether RADICaL simply ‘memorized’ the stereotyped trajectories for a limited number of conditions, or whether it could generalize to cases where each trial was more unique. To answer this question, we designed a “zero condition” simulation experiment, where each trial had its own unique Lorenz initial state and there were no repeated trials with the same underlying latent trajectories. (a) Example true (*top left*) and estimated Lorenz trajectories by RADICaL (*top right*), AutoLFADS (*bottom left*), and s-deconv (*bottom right*). Each trajectory is an individual trial, colored by the location of the initial state of the true Lorenz trajectory. The initial states of the trials are indicated by the dots in the same colors as the trajectories. (b) Performance in estimating the Lorenz *Z* dimension as a function of Lorenz oscillation frequency was quantified by variance explained ($R^2$) for all 4 methods.

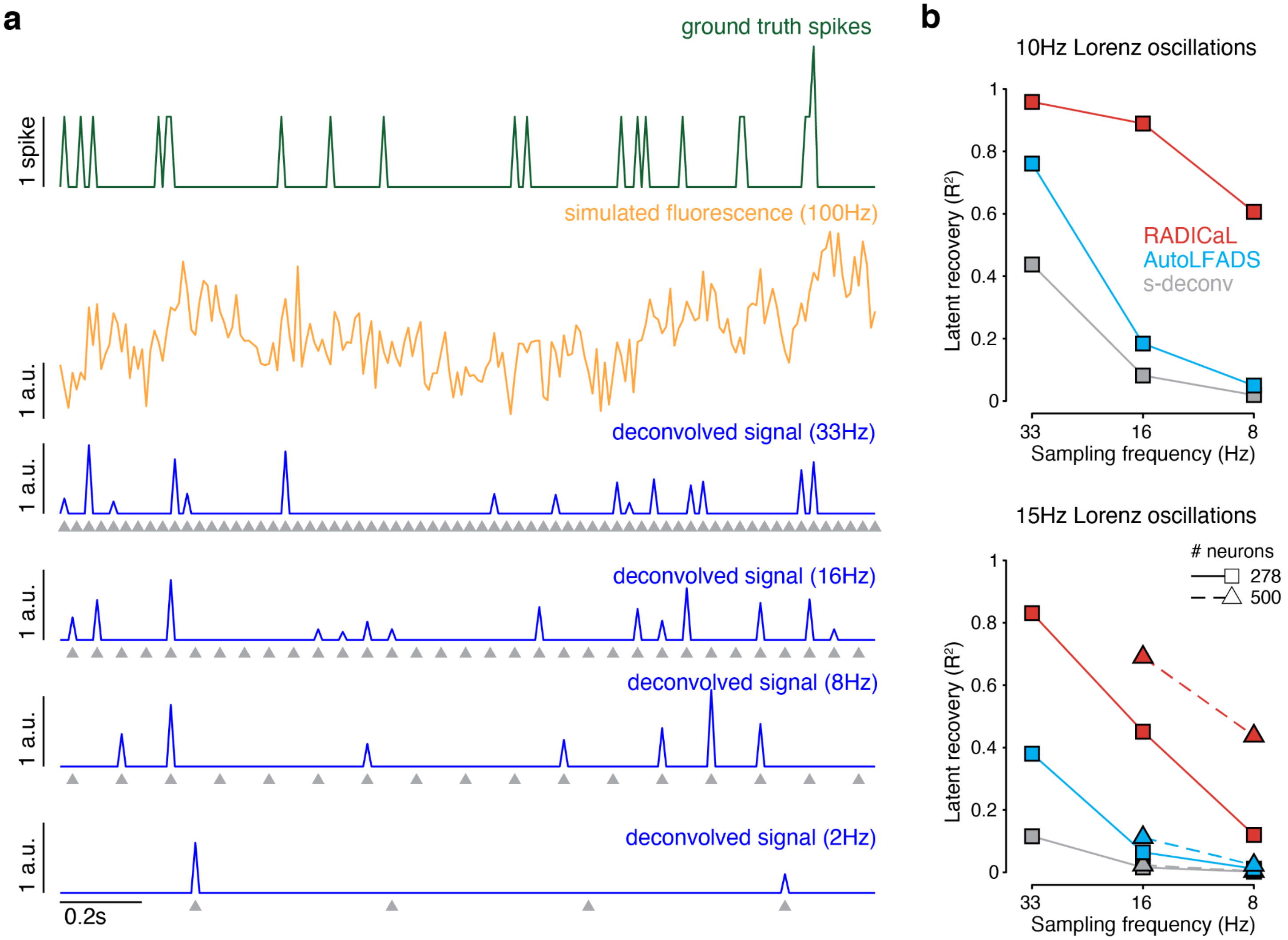


**Supplementary Figure 6 | RADICaL retains high latent recovery performance at slower imaging speeds, but there are limits to deconvolution with slower sampling.** To understand the extent to which the model performance depends on imaging speeds, we simulated data at different sampling rates ranging from 2 Hz to 33.3 Hz. (a) Example ground truth spikes, simulated fluorescence, and deconvolved signals at different sampling rates. Sample times are denoted by gray triangles. Deconvolution performance degraded at slower sampling rates, particularly in regimes when transients could be missed entirely. In our simulation we used a GCaMP6f model with a decay time of 400ms (see *Methods*). At an imaging rate of 2Hz, the majority of transients were missed and the estimate of the decay time constant tau was inaccurate (916.8 +/- 49.4ms, compared to the ground truth 400ms). Because deconvolution performs poorly at these sampling rates (i.e., <= 2Hz) with fast indicators, we do not recommend using RADICaL under such circumstances. (b) Performance in estimating the Lorenz *Z* dimension as a function of sampling rate was quantified by variance explained ($R^2$) for all 3 methods, for Lorenz oscillation frequencies of 10Hz (*top*) and 15Hz (*bottom*). Squares with solid lines denote experiments with 278 neurons. Triangles with dashed lines denote experiments with 500 neurons. RADICaL retained high performance and outperformed AutoLFADS and s-deconv in recovering the latent states of a 10 Hz Lorenz system at moderately slow sampling rates (8 and16 Hz; *top*). In real experiments, there may be benefits to slower sampling, e.g., one can image more neurons using a larger FOV. Increasing the number of neurons boosted RADICaL's performance, while AutoLFADS and s-deconv showed negligible improvement (*bottom*).

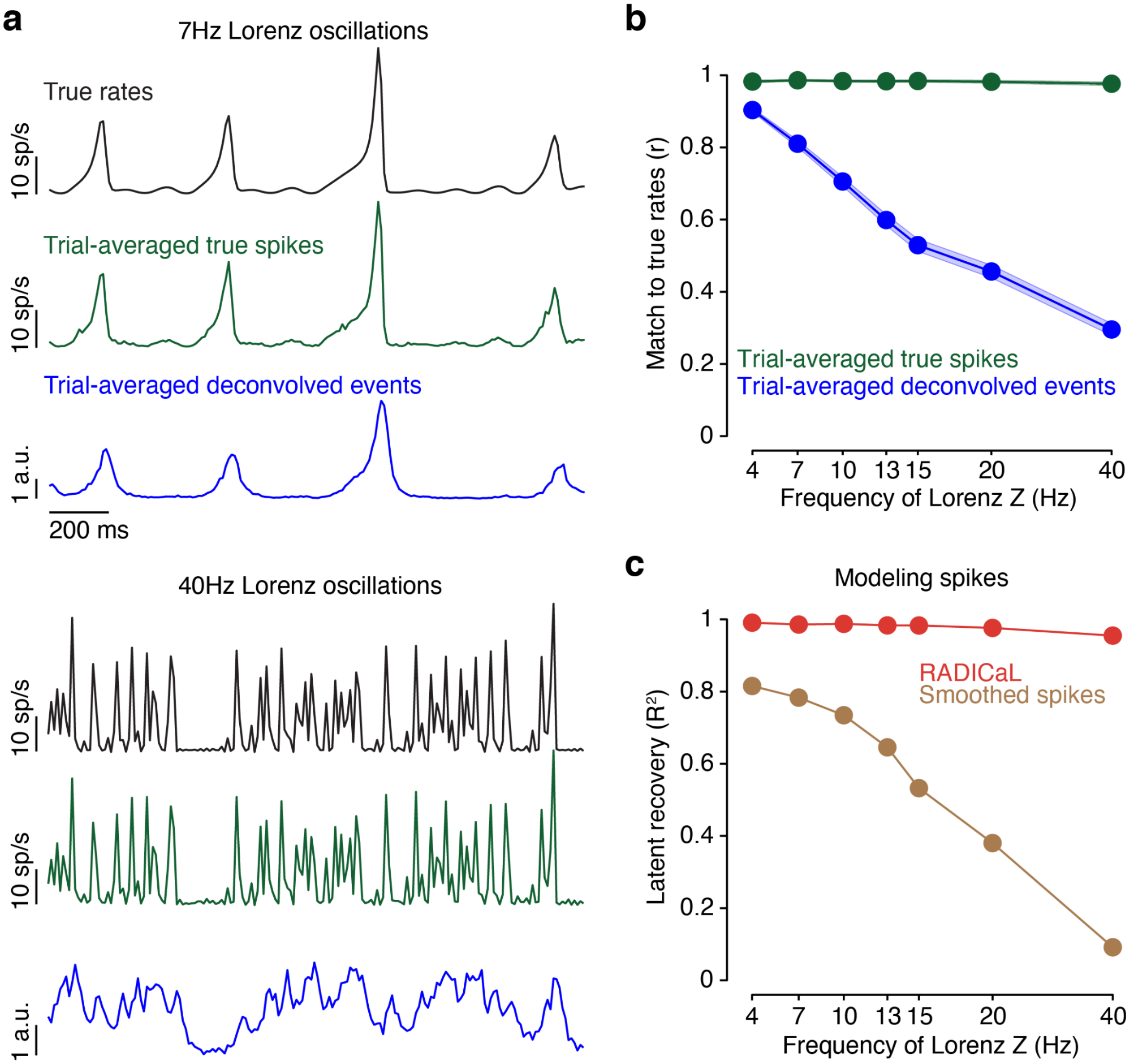


**Supplementary Figure 7 | Deconvolution places an upper bound on RADICaL's performance recovering higher-frequency features.** To understand how deconvolution performs across Lorenz oscillation frequencies, we measured how well trial-averaged deconvolved events captured the true underlying rate for individual (simulated) neurons. Averaging deconvolved events across the noisy repeated trials that have the same true underlying rates is a straightforward way to test, on average, whether deconvolution irreversibly loses information about the underlying rates. (a) Example ground truth firing rates, averaged true spikes across trials (3000 trials), and averaged deconvolved calcium events across trials (3000 trials), for Lorenz oscillation frequencies of 7Hz (*top*) and 40Hz (*bottom*). (b) Performance in capturing the ground truth firing rates as a function of Lorenz oscillation frequency was quantified by correlation coefficient (*r*) between the trial-averaged true spikes or deconvolved events and the true rates. Error bars indicate the variability across simulated neurons. The correlation between the trial-averaged deconvolved events and the true rate dropped as the Lorenz oscillation frequency increased, suggesting that deconvolution fails at higher Lorenz oscillation frequencies. (c) To determine whether RADICaL's performance loss for high-frequency signals was purely due to deconvolution failure or might involve limitations of the model itself, we eliminated the fluorescence generation/deconvolution step and applied RADICaL directly to the sub-sampled spiking activity. In this test, we did not use RADICAL's ZIG observation model, but kept the SBTT approach and used a Poisson observation model. Performance in using ground truth spikes to estimate the Lorenz *Z* dimension as a function of Lorenz oscillation frequency was quantified by variance explained ($R^2$) for smoothing and RADICaL. RADICaL retained high performance in latent recovery across Lorenz oscillation frequencies from 4Hz to 40Hz, whereas smoothing showed a much faster degradation of latent recovery performance. Together, these analyses demonstrate that the degradation in RADICaL's performance at higher Lorenz oscillation frequencies is mainly due to inaccuracies in deconvolution, and not due to the model itself.

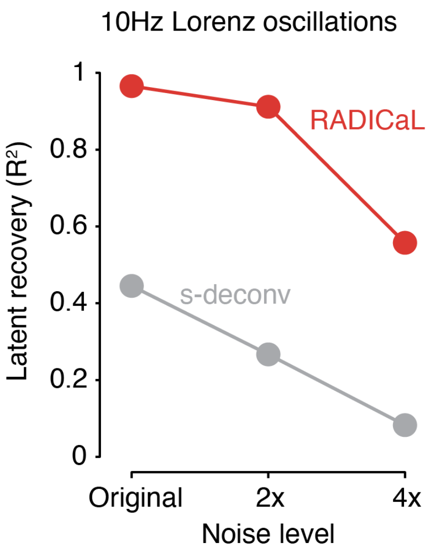


**Supplementary Figure 8 | Model tolerance to spike inference noise.** In our simulations, we chose parameters so that the resulting signal-to-noise regime produced similar correlations between real and inferred spike trains as observed in a recent benchmarking study[13] (see *Methods*). However, the spike inference noise can vary in real experiments and could affect RADICaL's performance. To test how larger spike inference noise affects the performance of RADICaL and s-deconv, we raised the level of the Gaussian noise used in generating simulated fluorescence traces by 2x or 4x. Performance in estimating the Lorenz *Z* dimension as a function of the level of spike inference noise was quantified by variance explained ($R^2$) for RADICaL and s-deconv. Performance declined for both methods as the noise level increased. However, RADICaL retained high performance at the 2x noise level ($R^2$=0.91) and reasonable performance at the 4x noise level ($R^2$=0.56). S-deconv had low performance across the board ($R^2$=0.27 and 0.08 for 2x and 4x noise, respectively). Notably, RADICaL performed better at the 4x noise level than s-deconv at the original noise level.

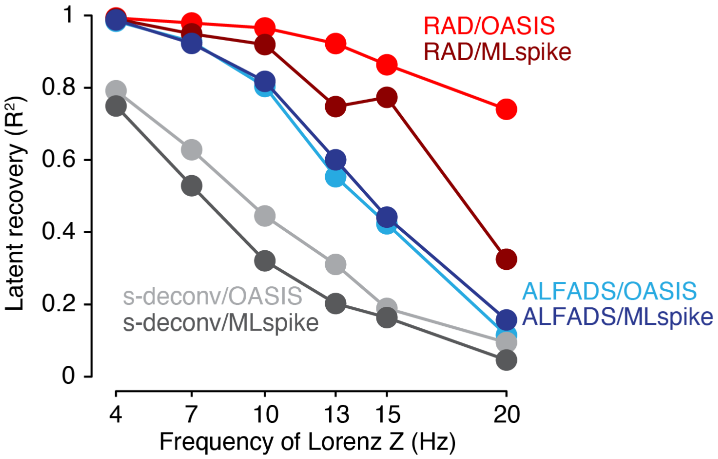


**Supplementary Figure 9 | RADICAL (with SBTT) improves latent recovery when using spikes inferred by MLspike, but does not perform as well as when using OASIS for deconvolution.** To test whether RADICaL could be effective with deconvolution algorithms that infer spike times instead of event rates, we analyzed simulated data that had spike inference performed with MLspike[27]. Performance in estimating the Lorenz *Z* dimension as a function of Lorenz oscillation frequency was quantified by variance explained ($R^2$) for six methods. These included three methods in which the inputs were deconvolved events from OASIS: RADICaL ("RAD/OASIS"), AutoLFADS ("ALFADS/OASIS") and smoothing ("s-deconv/OASIS"); and three methods in which inputs were spikes inferred with MLspike: RADICaL ("RAD/MLspike"), AutoLFADS ("ALFADS/MLspike") and smoothing ("s-deconv/MLspike"). When pairing RADICaL with deconvolution methods that produce spike times as output, we can use a Poisson observation model (as one would use for spikes measured via electrophysiology) instead of ZIG, while retaining the SBTT approach for sub-frame sampling. RADICaL with a Poisson observation model (RAD/MLspike) was able to model MLspike output, and substantially outperformed AutoLFADS and s-deconv (ALFADS/MLspike and s-deconv/MLspike), but did not perform as well as RADICaL applied to OASIS-deconvolved events (RAD/OASIS). In addition, the parameter tuning required for MLspike is more involved and requires more expertise than OASIS (see *Methods*). Therefore, we recommend using OASIS as the deconvolution method for RADICaL.

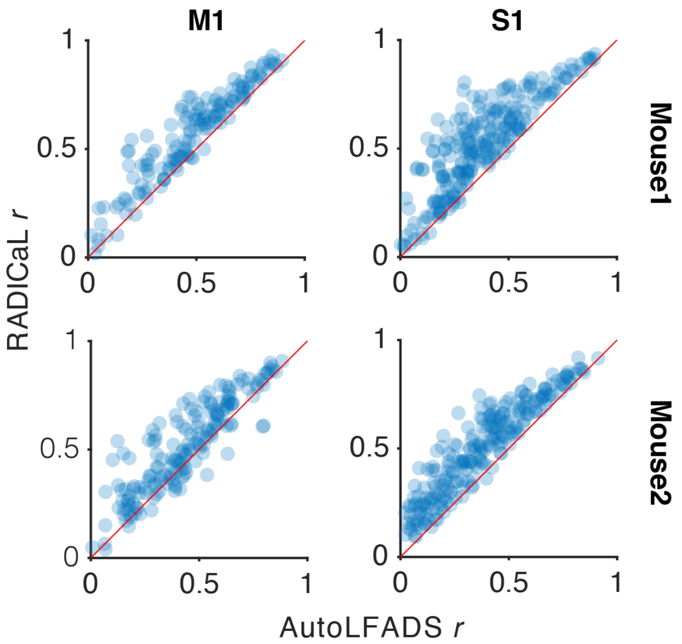


**Supplementary Figure 10 | Performance of RADICaL and AutoLFADS in capturing the empirical PSTHs on single trials in the mouse water grab experiments.** This figure is related to Figure 3d, but compares RADICaL with AutoLFADS instead of s-deconv. Correlation coefficient *r* was computed between the inferred single-trial event rates and empirical PSTHs. Each point represents an individual neuron. These results demonstrate that RADICaL captures the key features of individual neurons' responses from single-trial activity better than AutoLFADS in nearly every case.

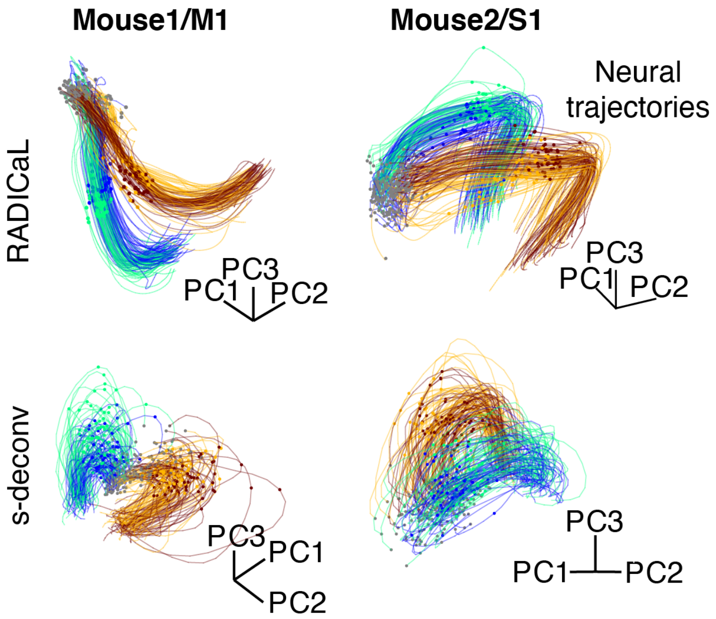


**Supplementary Figure 11 | Single-trial neural trajectories for additional mouse water grab experiments.** This figure is related to Figure 3e, and shows the remaining datasets. Single-trial, log-transformed event rates were projected into a subspace computed by applying PCA to the trial-averaged, log-transformed rates, colored by subgroups. Lift onset times are indicated by the dots in the same colors as the trajectories. Gray dots indicate 200 ms prior to lift onset time. *Top row*: single-trial neural trajectories derived from RADICaL rates; *Bottom row*: single-trial neural trajectories derived from s-deconv rates.

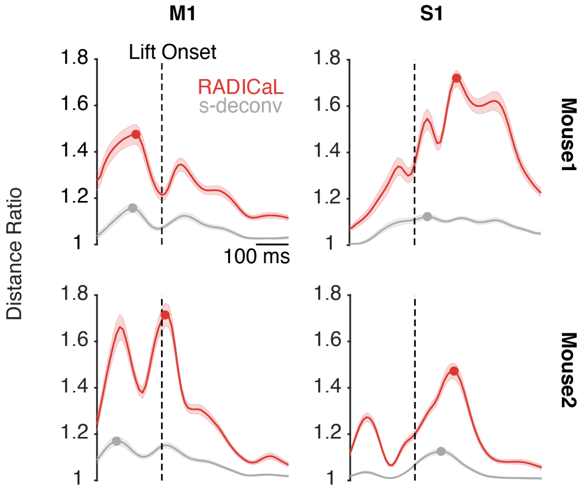


**Supplementary Figure 12 | RADICaL makes trial subgroups more distinct in an unsupervised manner.** To quantify how well trial subgroups were separated, we quantified the ratio of the cross-group distance to the within-group distance for each individual time point in a window from 200 ms before to 400 ms after lift onset (see *Methods*). A value larger than unity therefore indicates that the distance of trial in subgroup A to a trial in subgroup B is on average larger than the distance between trials of the same subgroup. This measure was computed separately for neural trajectories derived from RADICaL rates and from s-deconv rates. Horizontal scale bar represents 200 ms. Vertical dashed line denotes lift onset time. Error bar indicates the s.e.m. across individual trials. Dots indicate the maximum ratio for each method. The consistently larger values for RADICaL demonstrate that RADICaL produced neural trajectories that better reflected subgroup identity, despite being given no information about the subgroup membership or the kinematics used to define the subgroups.

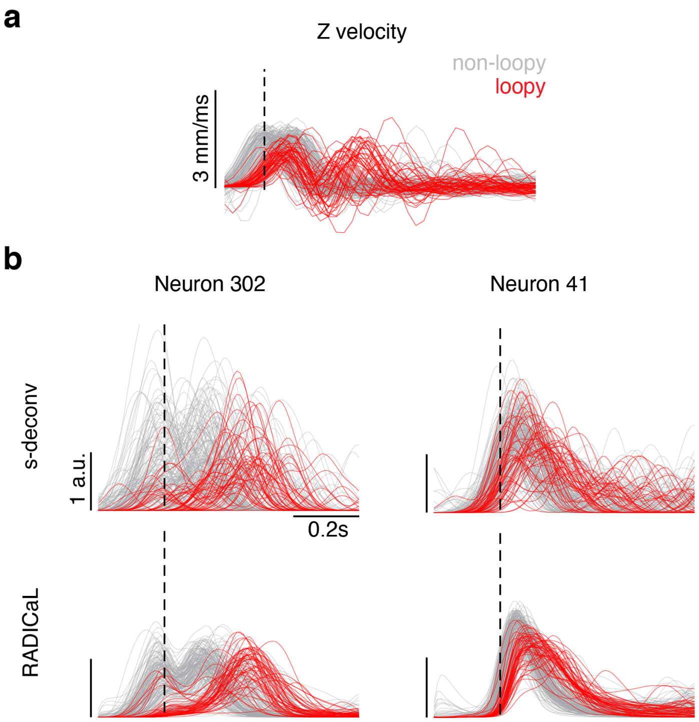


**Supplementary Figure 13 | RADICaL reveals distinct neural representations of "loopy" trials.** To determine whether RADICaL correctly captures large deviations from typical, we examined trials with unusual "loopy" paw kinematics. (a) Z-dimension of hand velocity profile. Each trace represents an individual trial, colored by loopy vs. non-loopy. Loopy trials are identified as the trials that have a second peak in Z-dimension of the hand velocity that is larger than 50% of the first peak. (b) Comparison of single-trial rates for 2 example neurons (data from Mouse1/S1) for s-deconv (*top row*) and RADICaL (*bottom row*). Each trace represents an individual trial (same color scheme as panel a). Horizontal scale bar represents 200 ms. Vertical scale bar denotes event rate (a.u.). Vertical dashed line denotes lift onset time. Note that the inferred single-trial firing rates for these neurons exhibited time courses that differed for loopy and non-loopy trials.

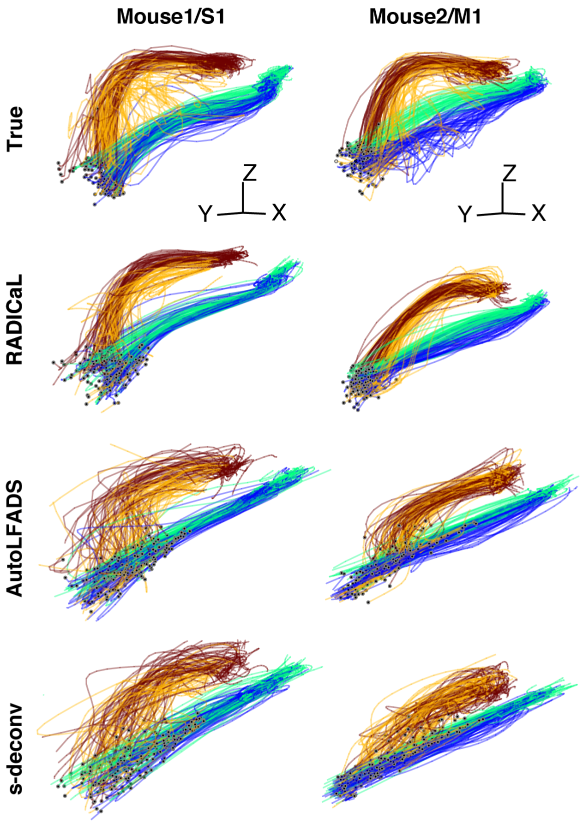


**Supplementary Figure 14 | Hand trajectories for additional mouse water grab experiments.** This figure is related to Figure 4a, and shows the remaining datasets. True and decoded hand positions for Mouse1/S1 (left) and Mouse2/M1 (right).

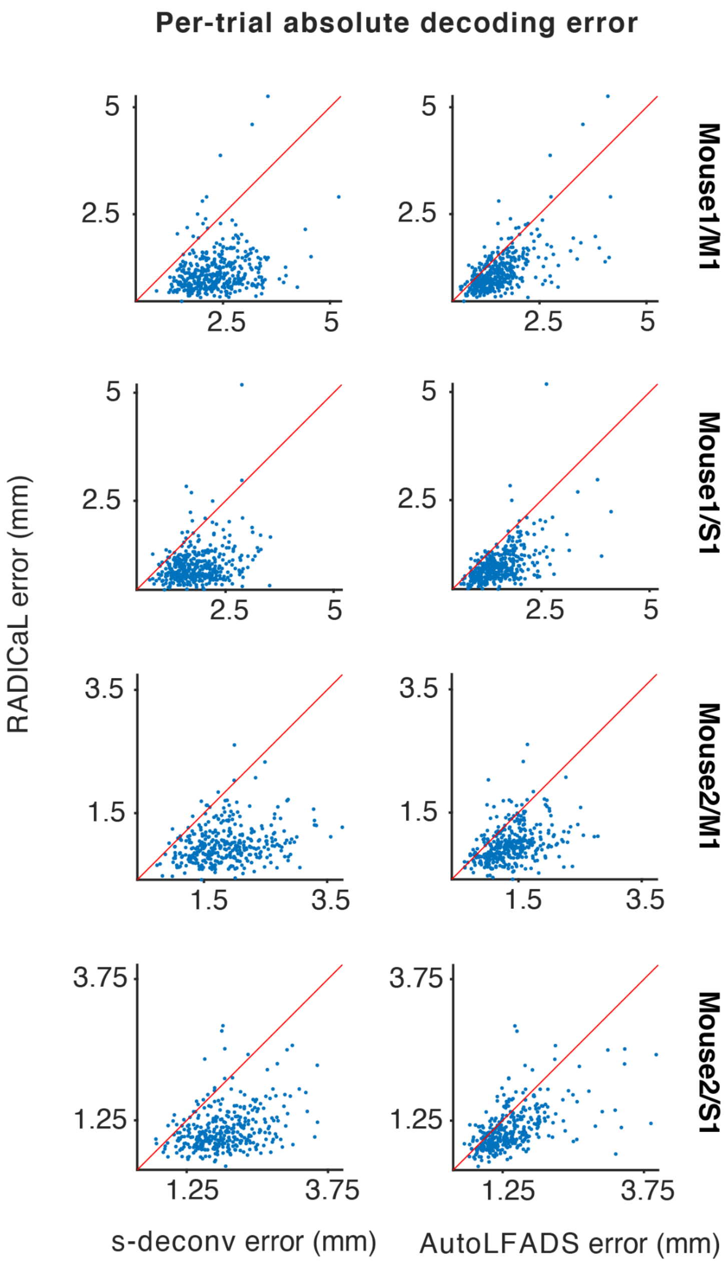


**Supplementary Figure 15 | RADICaL reduces decoding errors on the vast majority of single trials for all datasets.** Single-trial decoding error was quantified by measuring the absolute difference between the true and decoded hand position for each individual trial. Each point represents an individual trial. Error was greatly reduced compared with both s-deconv (*left*) and AutoLFADS (*right*).

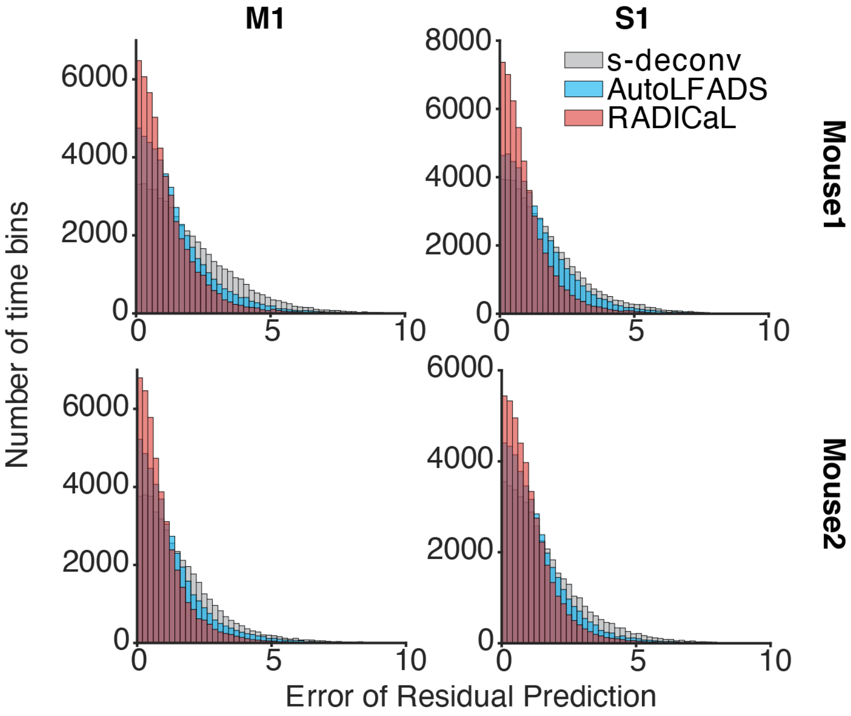


**Supplementary Figure 16 | RADICaL improves prediction of single-trial deviations from the mean of hand positions.** This figure demonstrates that RADICaL does not simply learn a ‘typical’ trajectory for left-reach trials and another for right-reach trials, but instead reflects small deviations from the condition average better than other methods. The residuals of hand positions (i.e., single-trial deviations from the mean) were computed by subtracting the left-reach or right-reach trial-averaged hand positions from the single trials. Error of residual prediction was computed by taking the absolute value of the difference between true and predicted residuals of hand positions.

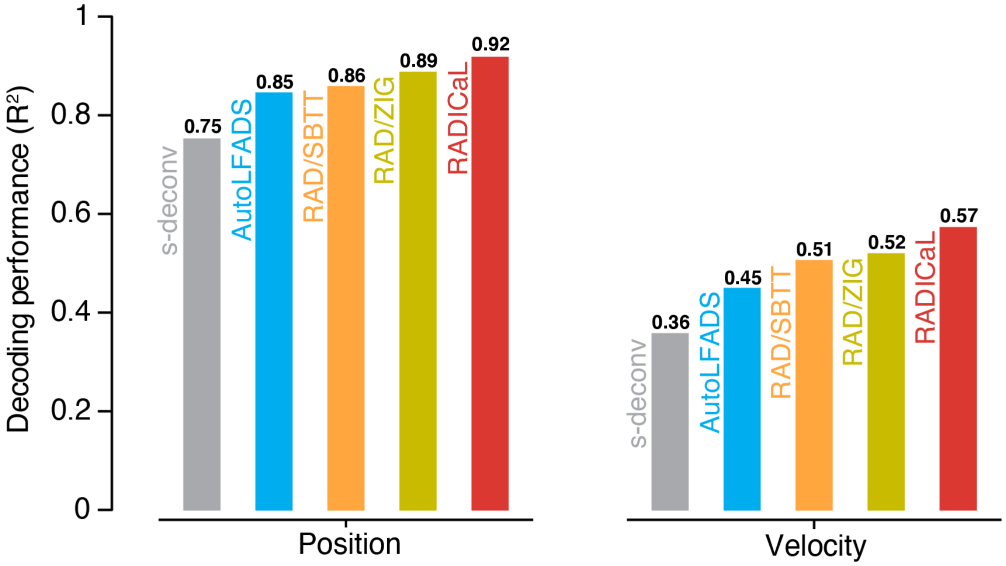


**Supplementary Figure 17 | Performance of ZIG-only (A-ZIG) and SBTT-only (A-SBTT) on decoding hand kinematics.** To test whether the innovations of RADICaL contributed separately to the improved decoding performance, we performed an ablation study where we enabled solely the ZIG emissions model (RAD/ZIG) or SBTT (RAD/SBTT). Decoding accuracy was quantified by measuring variance explained ($R^2$) between the true and decoded position (*left*) and velocity (*right*) across all trials, for RAD/ZIG, RAD/SBTT and other techniques. Analyzed data are from Mouse2/M1.

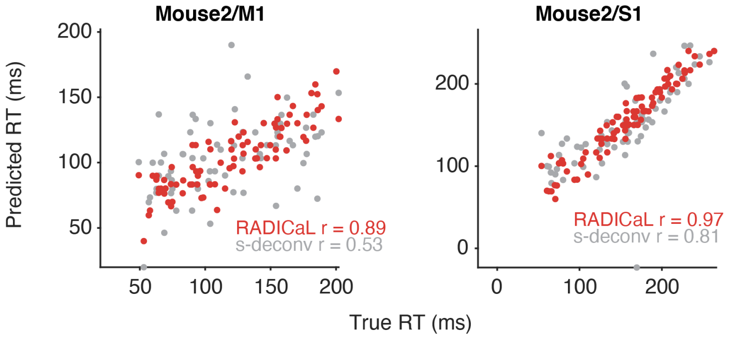


**Supplementary Figure 18 | Prediction of single-trial reaction times for additional mouse water grab experiments.** This figure is like Figure 4d, for the remaining datasets**.** Each dot represents an individual trial, color-coded by the technique. Correlation coefficient *r* was computed between the true and predicted reaction times. Data from Mouse2/M1 (*left*) and Mouse2/S1 (*right*).

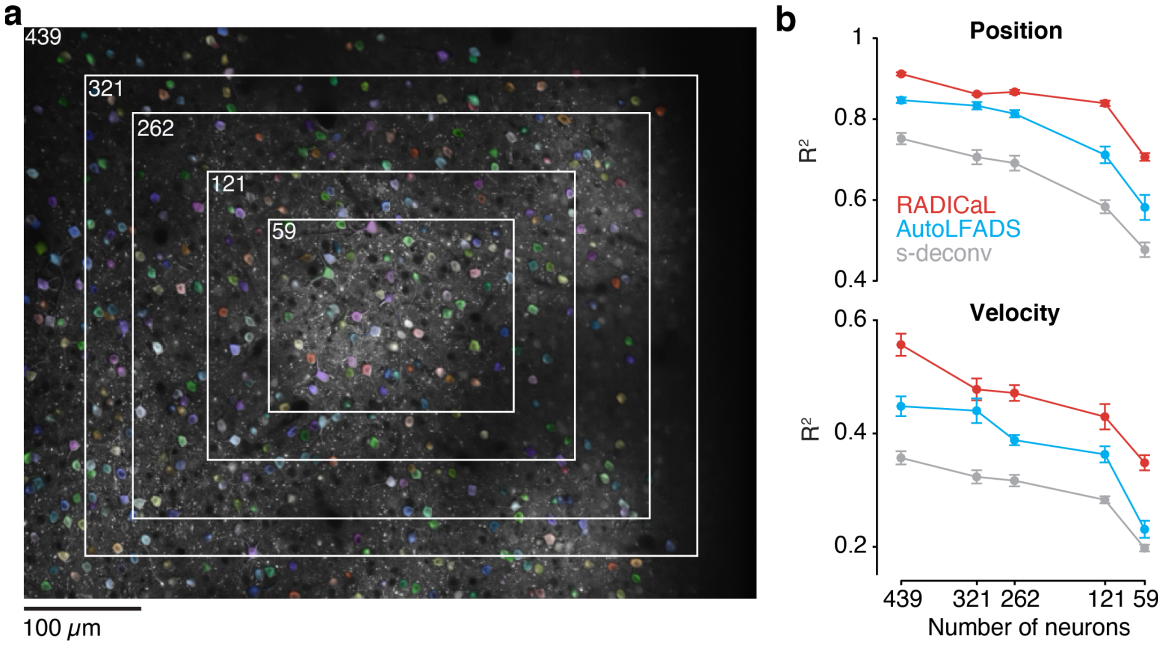


**Supplementary Figure 19 | RADICaL retains high decoding performance in an FOV-shrinking experiment.** This is an alternative method for evaluating performance with reduced neuron counts to the method in Figure 5. (a) The area selected to include was gradually shrunk to the center of the FOV to reduce the number of neurons included in training RADICaL or AutoLFADS. (b) Decoding performance measured using variance explained ($R^2$) as a function of the number of neurons used in each technique (top: Position; bottom: Velocity). Data from Mouse2/M1.

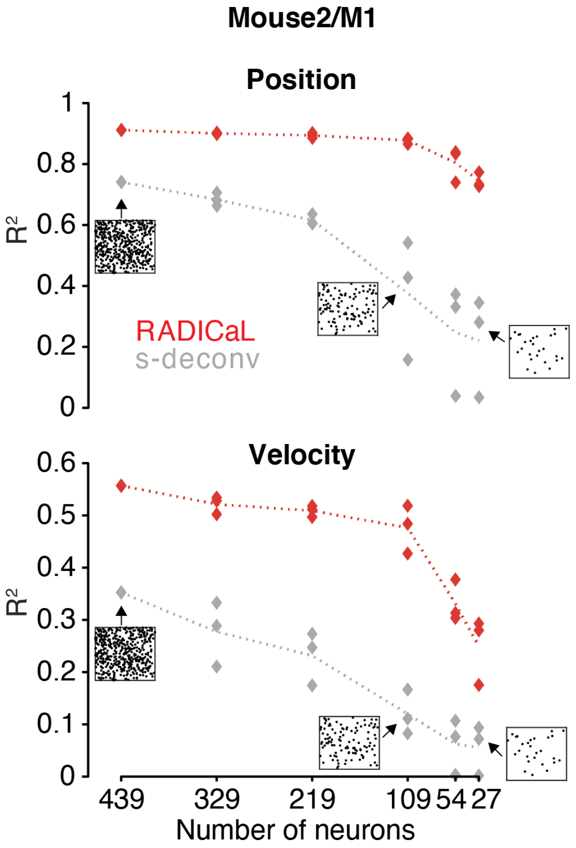


**Supplementary Figure 20 | RADICaL is robust to the random seed used in selecting subsets of neurons in a neuron downsampling experiment.** This figure is related to Figure 5. Decoding performance measured using variance explained ($R^2$) as a function of the number of neurons used in each technique (*top*: Position; *bottom*: Velocity). For a given number of neurons (except the full population of 439 neurons), 3 random seeds were used, and each data point represents an individual random seed. The dotted line represents the mean performance across the three random seeds for each method. Data from Mouse2/M1. Figure insets indicate the selected neurons in the FOV for experiments of the full population and example subsets of the population.

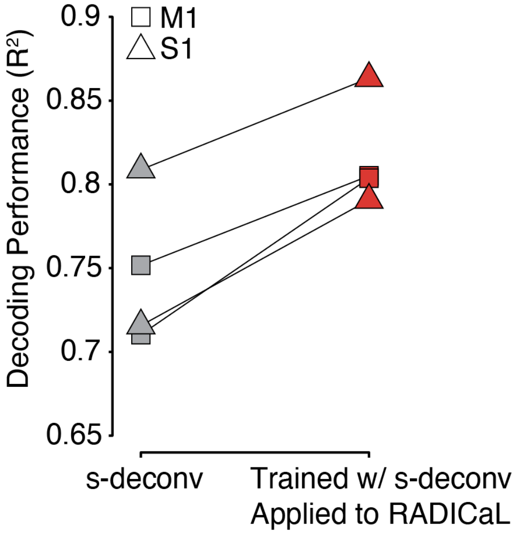


**Supplementary Figure 21 | RADICaL improves decoding performance using decoders trained with s-deconv rates.** This analysis demonstrates that the decoding performance benefit due to RADICaL cannot be due to training a better decoder alone, but results from better denoising of the trajectories themselves. Decoding performance was quantified by measuring variance explained ($R^2$) between the true and decoded hand position across each of the 4 datasets (2 mice for M1, denoted by squares, and 2 mice for S1, denoted by triangles), for decoders trained with s-deconv rates and applied to s-deconv rates (gray) or applied to RADICaL rates (*red*).

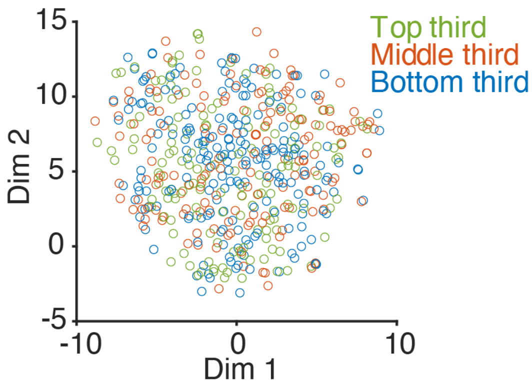


**Supplementary Figure 22 | Visualization of transformation from factors to neurons.** This analysis demonstrates that the different bands of the image use the same factors and not segregated ones, despite being divided up into separate sub-bins for improving temporal resolution with SBTT. The plot shows a 2-dimensional t-SNE space representation of weights mapping from RADICaL factors to ZIG parameters for Mouse1/M1. Each point represents an individual neuron (510 neurons total). Neurons are color coded based on the neurons' position within the field of view (i.e., top, middle, and bottom). The interspersal of the points shows that neurons do not have systematically different relationships with the factors in RADICAL based on which band they are in.
